# Biphasic Control of Cell Expansion by Auxin Coordinates Etiolated Seedling Development

**DOI:** 10.1101/2021.05.04.442657

**Authors:** Minmin Du, Firas Bou Daher, Yuanyuan Liu, Andrew Steward, Molly Tillmann, Xiaoyue Zhang, Jeh Haur Wong, Hong Ren, Jerry D. Cohen, Chuanyou Li, William M. Gray

## Abstract

Seedling emergence is critical for food security. It requires rapid hypocotyl elongation and apical hook formation, both of which are mediated by regulated cell expansion. How these events are coordinated in etiolated seedlings is unclear. Here, we show that biphasic control of cell expansion by the phytohormone auxin underlies this process. Shortly after germination, high auxin levels restrain elongation. This provides a temporal window for apical hook formation, involving a gravity-induced auxin maximum on the eventual concave side of the hook, triggering PP2C.D1controlled asymmetrical H^+^-ATPase activity, resulting in differential cell elongation. Subsequently, auxin concentrations decline acropetally and switch from restraining to promoting elongation, driving hypocotyl elongation. Our findings elucidate how differential auxin concentrations throughout the hypocotyl coordinate etiolated development, leading to successful soil emergence.

**One-Sentence Summary:** Auxin concentration-dependent cell expansion coordinates hypocotyl elongation and apical hook development for soil emergence.

## Main Text

Successful seedling emergence enables plants to produce photosynthates to fuel growth and development. In most dicots, rapid hypocotyl elongation drives seedling emergence from the soil(*1*). Simultaneously, the apical hook structure develops at the hypocotyl apex, which protects the shoot apical meristem from damage(*2, 3*). This strategy of seedling emergence was described 140 years ago in Darwin’s pioneering work(*2*). Auxin (indole-3-acetic acid, IAA) is indispensable for regulating both hypocotyl elongation and apical hook development(*4-9*), through its ability to mediate changes in cell expansion in a tissue- and concentration-dependent manner(*10*). However, key questions related to these processes remain. First, while auxin can promote cell elongation in the growing parts of the hypocotyl(*4*), it inhibits elongation at the apical hook region(*9*). How auxin exerts these differential effects throughout the hypocotyl is unknown. Second, auxin asymmetrically accumulates at the concave side of the hook to inhibit cell elongation and to drive hook formation(*5, 9*). However, the signal that initiates the asymmetric auxin distribution remains uncertain(*11, 12*). And third, auxin-mediated promotion of hypocotyl cell elongation can be explained by the acid growth theory(*13*). This requires auxin-induced Small Auxin Up RNA (SAUR)-mediated repression of PP2C.D phosphatases to activate plasma membrane (PM) H^+^ATPases(*4, 14-16*). These downstream effectors of auxin action may also participate in apical hook development(*15, 16*), but the mechanism of inhibition of cell elongation on the concave side of the hook has yet to be identified.

### Auxin regulates hypocotyl cell elongation in a biphasic manner

Auxin regulates cell expansion in a concentration- and tissue-dependent manner(*4*). In general, auxin promotes cell expansion in shoots(*15*) while inhibiting it in roots(*17, 18*). A notable exception exists at the concave side of the apical hook, where auxin inhibits cell expansion(*9*). To obtain insights into how auxin coordinates hypocotyl elongation and apical hook development, we first analyzed the spatial distribution of auxin signaling in the epidermis of etiolated *Arabidopsis* hypocotyls, in which cell elongation proceeds acropetally from the base to the tip over time(*1, 19*). We used the *R2D2* reporter, which consists of auxin-degradable (DII) and auxin-insensitive (mDII) fluorescent proteins(*20*). The mDII/DII signal ratio may serve as a proxy for cellular auxin levels(*20*). IAA treatment increased the mDII/DII ratio whereas the auxin biosynthesis inhibitor KOK2153(*21*) led to a decrease in the ratio (fig. S1), validating the use of this reporter as a proxy for auxin distribution in these cells. Quantification of the mDII/DII ratio suggested an auxin gradient along the hypocotyl, with a higher auxin signal in the top cells (Fig. 1, A and B, fig S1). This gradient was confirmed by IAA quantification in the upper and lower halves of hypocotyls (fig. S1D).

**Fig. 1.**
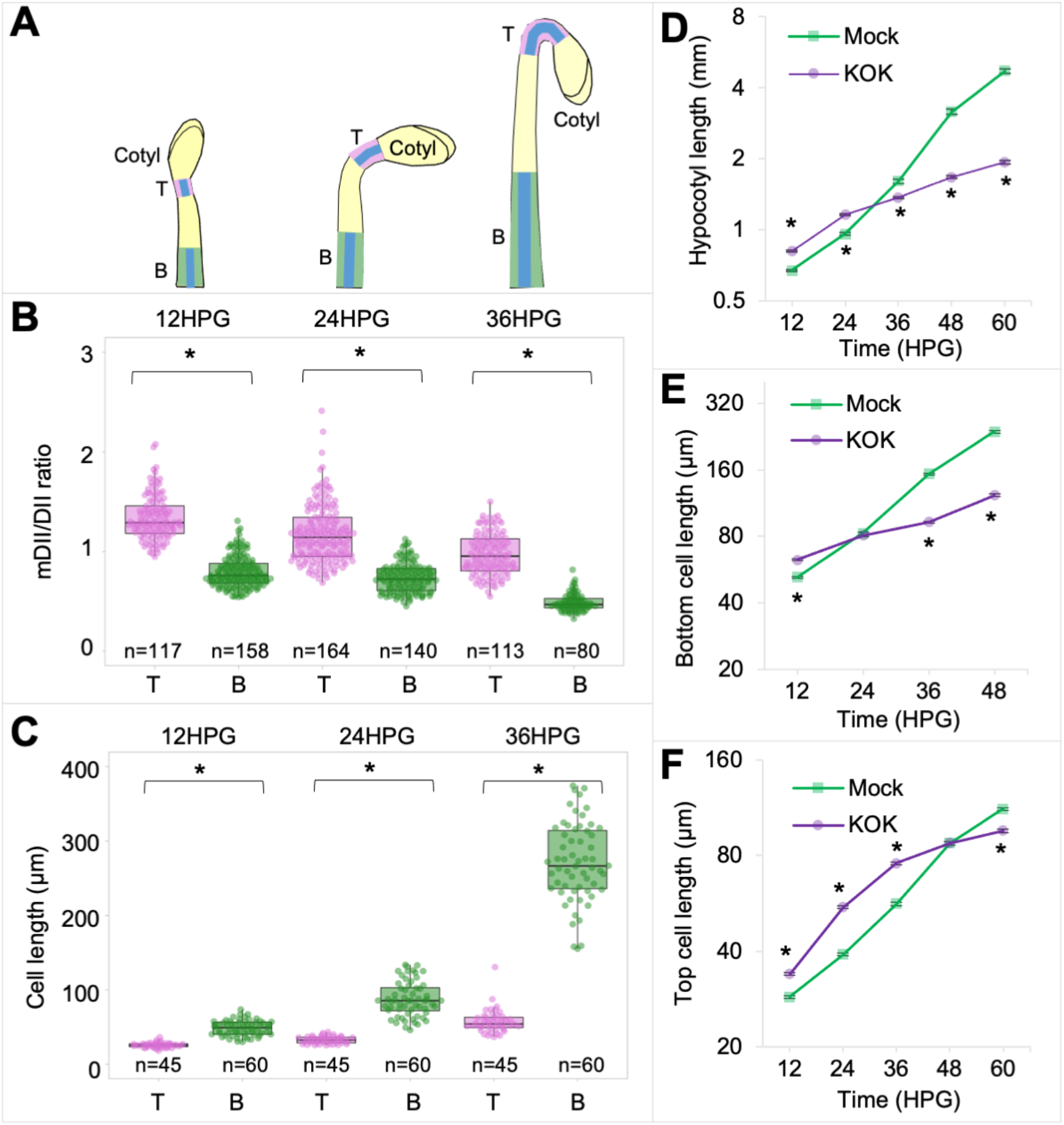
Auxin regulates hypocotyl cell elongation in a biphasic manner. **(A)** Schematic diagrams showing the positions of the top cells (T) and bottom cells (B) for analysis in etiolated hypocotyls. The epidermal cells where the *R2D2* signals and cell lengths were measured are highlighted in blue. (**B** and **C)** Quantification of the mDII/DII ratio (**B**) and cell lengths (**C**) in the top (T) and bottom (B) cells of *R2D2* hypocotyls for the indicated time. Twenty epidermal hypocotyl cells were numbered (No.1-20) from the collet to the cotyledons. Four cells (No. 2-5) from the bottom (B) and three cells (No. 18-20) from the top (T) were used for quantification. * indicates *P* value < 0.01. **(D)** Quantification of hypocotyl length during etiolated seedling development, n=60 hypocotyls. (**E** and **F)**, Quantification of epidermal cell length of the bottom (**E**) and top (**F**) cells during etiolated seedling development, n=300 cells. For (**D** to **F)**, germinated seeds were transferred to ½ MS medium containing 1% sucrose supplemented with DMSO (Mock) or KOK2153 (KOK). Five bottom cells (No. 2-6) and five top cells (No. 16-20) were used for quantification. Values represent sample means ± s.e.m. from three replicates. * indicates *P* value < 0.01.

An inverse correlation between auxin level and cell length in the hypocotyl (Fig. 1C) suggested a critical role for the auxin gradient in regulating hypocotyl elongation. We therefore undertook a detailed analysis of hypocotyl and cell lengths during early seedling development. Etiolated hypocotyls showed a transition from slow to rapid elongation beginning at around 24 hours post germination (HPG) (Fig. 1D). Inclusion of KOK2153 in the medium significantly promoted hypocotyl elongation at the early stages (before 24HPG) but inhibited elongation thereafter (Fig. 1D). Consistently, epidermal cell lengths in the presence of the inhibitor were significantly increased at the early stages but decreased at the later stages (Fig. 1, E and F), compared with untreated controls. The inhibitor’s effects were largely reversed by co-treatment with 100 nM IAA, indicating that these effects were caused by inhibition of auxin biosynthesis (fig. S2, A and B). These observations suggest a biphasic role for auxin in hypocotyl cell elongation: auxin inhibits cell elongation at the early stages but promotes it at the later stages. Consistent with this notion, treatment with auxinole, an auxin antagonist of the SCF^TIR1/AFB^ receptors(*22*), also promoted hypocotyl cell elongation at 12HPG (fig. S2, C to E). To genetically challenge this hypothesis, we analyzed the *A. thaliana* mutants *wei8-3 tar2-1* and *tir1 afb2* that display decreased auxin biosynthesis(*23, 24*) and defective auxin perception(*25*), respectively. Both mutants exhibited longer hypocotyl cells at 12HPG when compared with the wild type (WT) (fig. S2, F to H). By contrast, the *yuc1-D* mutant, which overproduces IAA(*26*), displayed shorter hypocotyl cells (fig. S2, F to H). Although auxin is typically thought to promote cell elongation in shoots, here we demonstrate, through fine temporal and spatial analyses, an inhibitory effect on hypocotyl cell elongation during the early stages of etiolated seedling development.

### Auxin concentrations correlate with the biphasic hypocotyl elongation

The biphasic role of auxin in hypocotyl cell elongation indicates a transition in auxin’s role from restraining to promoting growth. This transition occurred earlier in the bottom cells (at ∼24HPG) than in the top cells (after 36HPG) (Fig. 1, E and F). Interestingly, the cell length at which the transition occurred was very similar between the bottom and top cells (∼80 µm; Fig. 1, E and F), suggesting that cell size might be a key factor affecting the transition. Because the R2D2 reporter suggested that bottom cells contained less auxin (Fig. 1B and fig. S1D), we hypothesized that the transition in auxin’s role from restraining to promoting cell elongation is driven by the decreased IAA concentration in cells to levels that promote elongation(*10*). In fact, as hypocotyls elongated, the mDII/DII ratio in both bottom cells and top cells gradually dropped, and the ratio at which the transition occurred was very similar between the bottom (24 HPG) and top (36HPG) cells (∼0.8; fig. S3, A and B). These findings support the idea that the transition is the result of reduced cellular IAA concentration. Conceivably, this reduction in IAA concentration could be caused by the slow increase in cell volume that occurs during the time at which auxin is restraining elongation. To explore this possibility, we quantified the IAA concentration in elongating hypocotyls over time and found a significant decrease in auxin concentration (as expressed per volume of hypocotyl) between 12, 24 and 36HPG (fig. S3, C and D).

We next examined how this decrease in auxin concentrations is reflected on the expression of *SAUR* genes, which promote auxin-mediated cell elongation. The expression of *SAUR* genes increased as hypocotyls grew (fig. S3, E and F), suggesting that high IAA levels are inhibitory on *SAUR* expression at the early stages of hypocotyl elongation. Consistent with this possibility, 1 µM IAA treatment repressed the expression of *SAUR* genes but induced the expression of other auxin-responsive genes in hypocotyls (fig. S3G). These results suggested that auxin-mediated repression of *SAUR* expression might contribute to its inhibitory effect on hypocotyl cell elongation at the early stages. This notion is supported by the observation that constitutive expression of *SAUR19* from the 35S promoter resulted in increased hypocotyl cell length at 12HPG (fig S3, H to J).

Together, these findings suggest that high auxin levels inhibit *SAUR* expression and cell elongation during the early stages of etiolated seedling development. As the hypocotyl cells elongate following an acropetal wave, the auxin concentration drops, perhaps as a result of the increased cell volume, and auxin switches to promote cell elongation. This concentrationdependent inhibition suggests parallels with apical hook formation, where high IAA levels in cells of the presumptive concave side of the developing hook are believed to inhibit cell elongation(*6*). We therefore undertook an investigation to determine whether and how this inhibition is involved in apical hook formation during early seedling development.

### Gravity triggers asymmetric auxin distribution for apical hook formation

Apical hook formation is driven by asymmetric auxin distribution across the hypocotyl(*3, 5, 27*). The signal that initiates the asymmetric auxin distribution remains unclear(*11, 12*). Gravity has long been suggested to play a role in apical hook formation(*28-30*), and recently a signal originating from gravity-induced root bending, but not from gravity *per se*, was proposed to direct apical hook formation(*11*). In that study, apical hooks still formed after removal of the root(*11*), suggesting the signal may not originate from the root. We therefore hypothesized that apical hook formation is a hypocotyl gravitropic response. We first noticed that, upon rotating etiolated seedlings to a horizontal plane, the hypocotyl curved against the gravity, while apical hooks actively reoriented positively with the new gravity vector (movie S1). Because shoot gravitropic response was reported to be sensitive to the inclination angle of the organ from the direction of gravity(*31, 32*), we tested whether the inclination angle of the hypocotyl would affect hook formation. We placed the germinated seeds at a vertical orientation (inclination angle *θ* = 0, vertical group) or at a horizontal orientation (inclination angle *θ* = 90°, horizontal group) (Fig. 2A and fig. S4A) and monitored the development of the hook. Intriguingly, apical hooks formed earlier in the horizontal group than in the vertical group, regardless of cotyledon orientation (Fig. 2A, fig. S4A and movie S2). In the horizontal group, hook bending was observed as early as 6HPG, and the hook angle reached its maximum around 24HPG. By contrast, in the vertical group, hypocotyls started to bend at about 12HPG and fully formed the hook after 30HPG. The asymmetric distribution of auxin signal, as shown by the *DR5::VENUS-NLS* reporter(*33*), was also established earlier in the horizontal group than in the vertical group (Fig. 2B). Furthermore, hook formation was affected by counteracting the effect of gravity via periodically rotating the seedlings (fig. 4, B and C). These data indicate that hook formation is a gravity-dependent response. To genetically support this notion, we analyzed mutants that are defective in shoot gravitropism. Two radial pattern mutants, *scarecrow* (*scr-3*/*shoot gravitropism1*) and *short-root* (*shr-2*/*shoot gravitropism7*), exhibit abnormal shoot gravitropism, due to the absence of the gravity-sensing endodermal cell layer(*34*). Both mutants were defective in apical hook formation, with the more severe *shr-2* mutant exhibiting a stronger defect (Fig. 2C and movie S3). Consistently, the normally asymmetric distribution of the *DR5::VENUS-NLS* signal at the hook region was largely abolished in the *shr-2* mutant (Fig. 2D). Recently, the *LAZY* family of genes have been implicated in the control of gravity-induced directional auxin transport(*35*). In the *atlazy2,3,4* triple mutant and the *atlazy1,2,3,4* quadruple mutant, auxin accumulated at the non-gravistimulated side and, accordingly, the mutants exhibited reversed gravitropic responses(*36*). Hence, we employed these *atlazy* mutants to examine gravity’s role in apical hook formation. When placed at a horizontal orientation, 14% of the *atlazy2,3,4* hooks (8/57) and 92% of the *atlazy1,2,3,4* hooks (56/60) were formed with the concave side facing upwards instead of downwards (Fig. 2, E and F; movie S4), and DR5::GFP(*37*) signal accumulated at the non-gravistimulated side (Fig. 2G). Furthermore, consistent with a previous study showing that gravity-induced PIN3 polarization mediated auxin flow toward the lower side of hypocotyl(*38*), we found that, when the hook is forming, PIN3GFP(*8*) had a stronger accumulation in the endodermis of the concave side compared to the convex side (Fig. 2H), supporting that gravity-induced PIN3 polarization is also involved in auxin mobilization for apical hook formation. These results strongly support that gravity is the initial signal that triggers asymmetric auxin distribution for apical hook formation, and suggest it is a gravitropic response of the young hypocotyl.

**Fig. 2.**
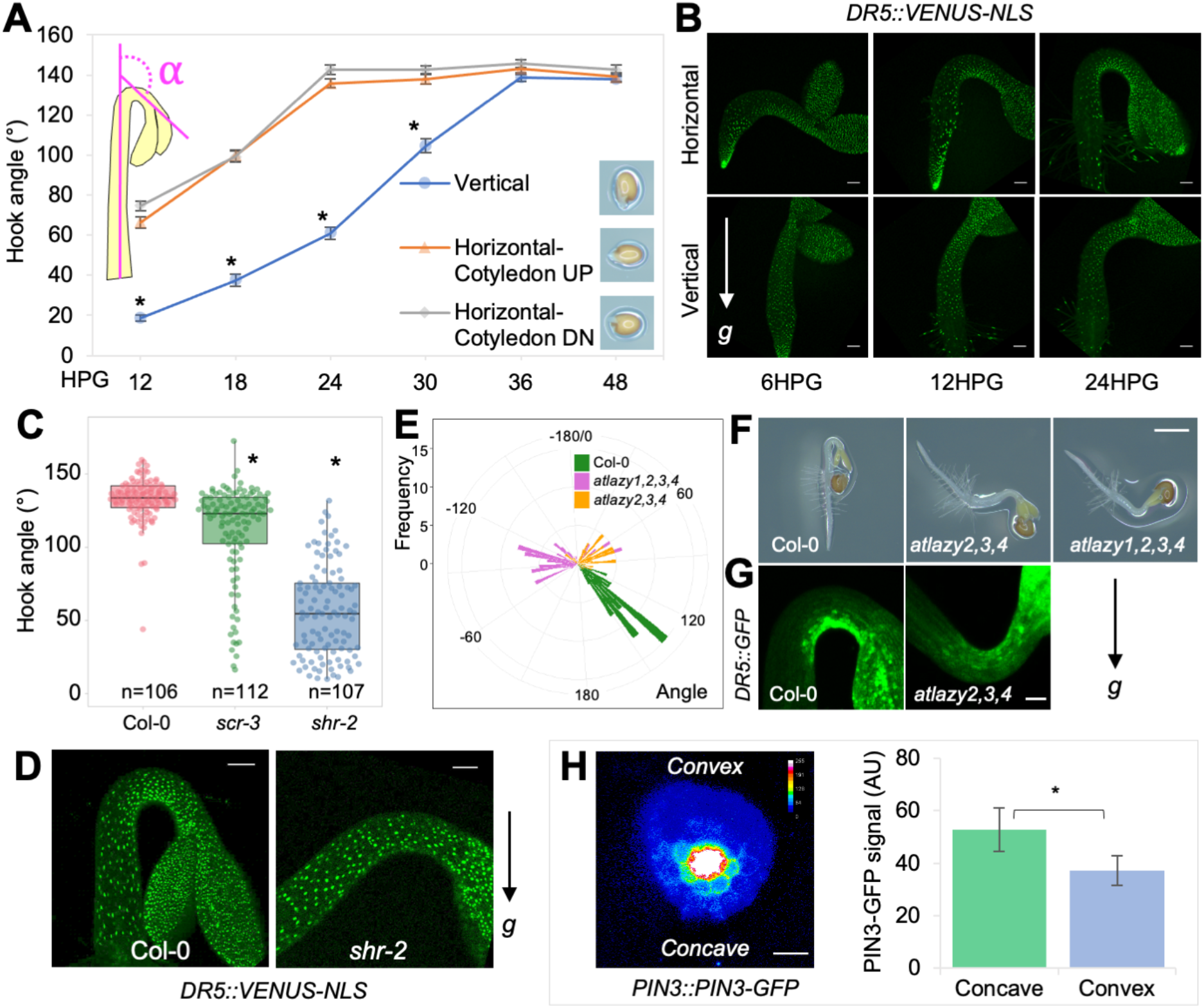
Gravity triggers asymmetric auxin distribution during apical hook formation. **(A)** Quantification of the kinetics of apical hook development in differentially oriented seedlings. Germinated seeds were initially placed at a vertical orientation or horizontal orientations, for which cotyledons were oriented either above or below of the hypocotyl. Hook curvature angles were measured over 48 h; n=36 hooks. Values represent sample means ± s.e.m. from three replicates. The inset depicts how the angle of hook curvature was determined. **(B)**, *DR5::VENUS-NLS* expression during apical hook development in differentially oriented seedlings. Scale bars=100 µm. (**C)** Quantification of the hook angle of Col-0 and *scr-3* and *shr-2* mutants. (**D)** *DR5::VENUS-NLS* expression in the apical hook of Col-0 and *shr-2*. Scale bars=100 µm. **(E)** Quantification of the hook angle and direction of Col-0 and *atlazy2,3,4* and *atlazy1,2,3,4* mutants. Hook curvature angles of concave-side-up seedlings have negative values. **(F)** Representative pictures showing the apical hook of Col-0, *atlazy2,3,4* and *atlazy1,2,3,4*. Scale bar=1 mm. **(G)** *DR5::GFP* expression in the apical hook of Col-0 and an *atlazy2,3,4* seedling that developed a concave-side-up hook. Scale bar=100 µm. For **(C** to **G)**, germinated seeds were initially placed at a horizontal orientation and the hook pictures were acquired at 24HPG. **(H)** Subcellular localization of PIN3-GFP in a transverse section of the hook at 6HPG. Scale bar=50 µm. Description of the quantification of the PIN3-GFP signals in the endodermis of the concave and convex side can be found in the methods section. For (**A, C** and **H)**, * indicates *P* value < 0.01.

**Fig. 3.**
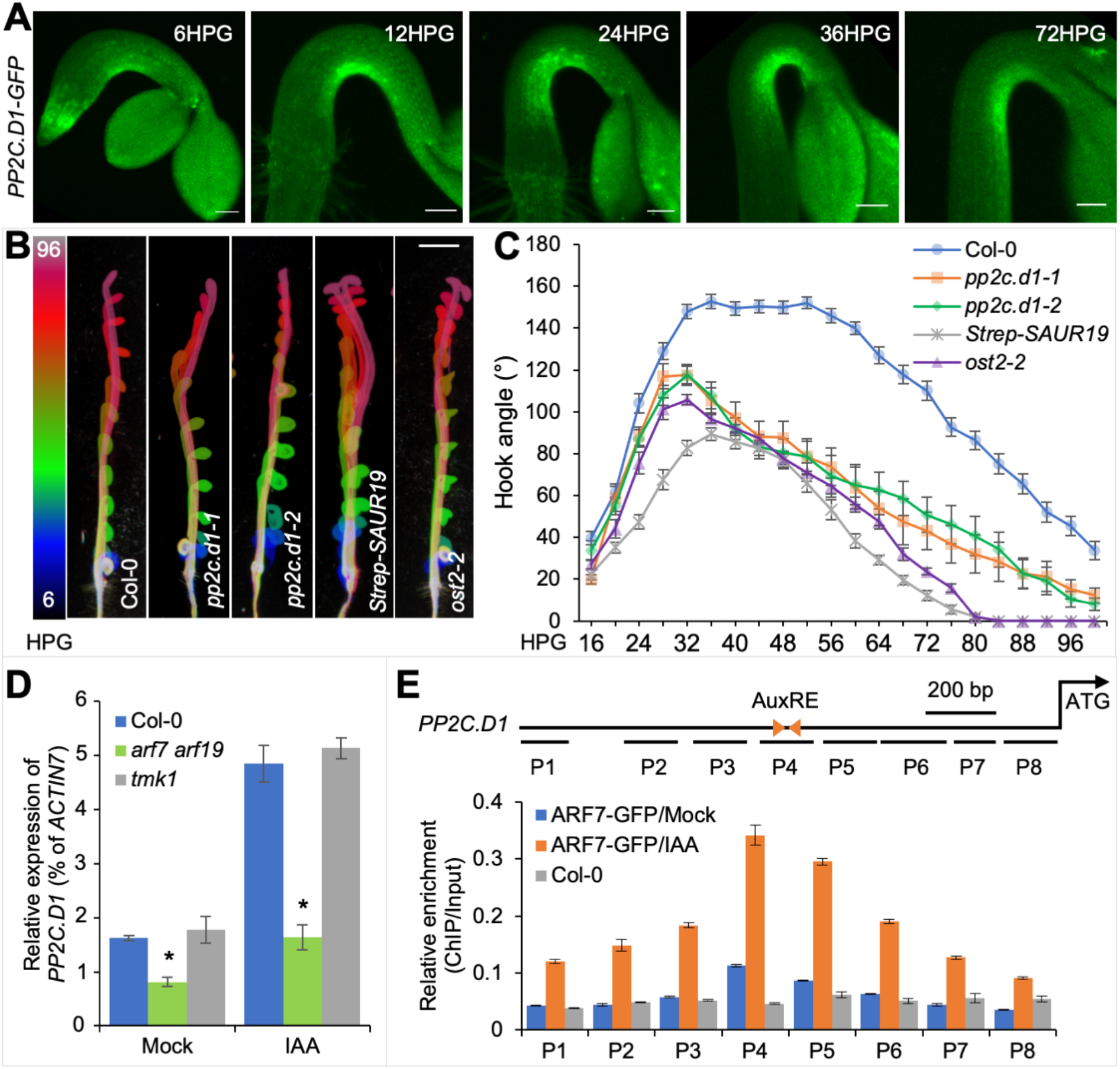
Auxin induces *PP2C*.*D1* expression via ARF7. **(A)** *PP2C*.*D1::PP2C*.*D1-GFP* expression during apical hook development. Germinated seeds were initially placed at a horizontal orientation at 0HPG. Scale bars=100 µm. **(B)** Apical hook development in Col-0, *pp2c*.*d1-1, pp2c*.*d1-2, 35S::Strep-SAUR19* and *ost2-2*. The color scale indicates the time points (6-96HPG) with 6-hour intervals. Scale bar=2 mm. **(C)** Kinetics of apical hook development in Col-0 (n=39), *pp2c*.*d1-1* (n=13), *pp2c*.*d1-2* (n=16), *35S::Strep-SAUR19* (n=22) and *ost2-2* (n=25). Values represent sample means ± s.e.m. from three replicates. **(D)** RT-qPCR analysis of *PP2C*.*D1* expression in the whole seedlings of WT, *tmk1-1*, and *arf7 arf719* double mutant. Three-day-old dark-grown seedlings were transferred to 1/2 MS liquid medium containing 1% sucrose with or without 10 µM IAA for 2 hours. *PP2C*.*D1* transcript levels were normalized against *ACTIN7* expression. Data are means ± s.e.m. from three biological replicates. * indicates *P* value < 0.01. **(E)** ChIP-qPCR assays showing the binding of ARF7-GFP to the *PP2C*.*D1* promoter. Schematic representation of the *PP2C*.*D1* gene. Triangles represent the inverted AuxREs and short lines represent the DNA probes followed in the ChIP-qPCR assays. Each ChIP value was normalized to its respective input DNA value, and enrichment is shown as the percentage of input. Error bars represent the s.e.m. of three biological replicates.

**Fig. 4.**
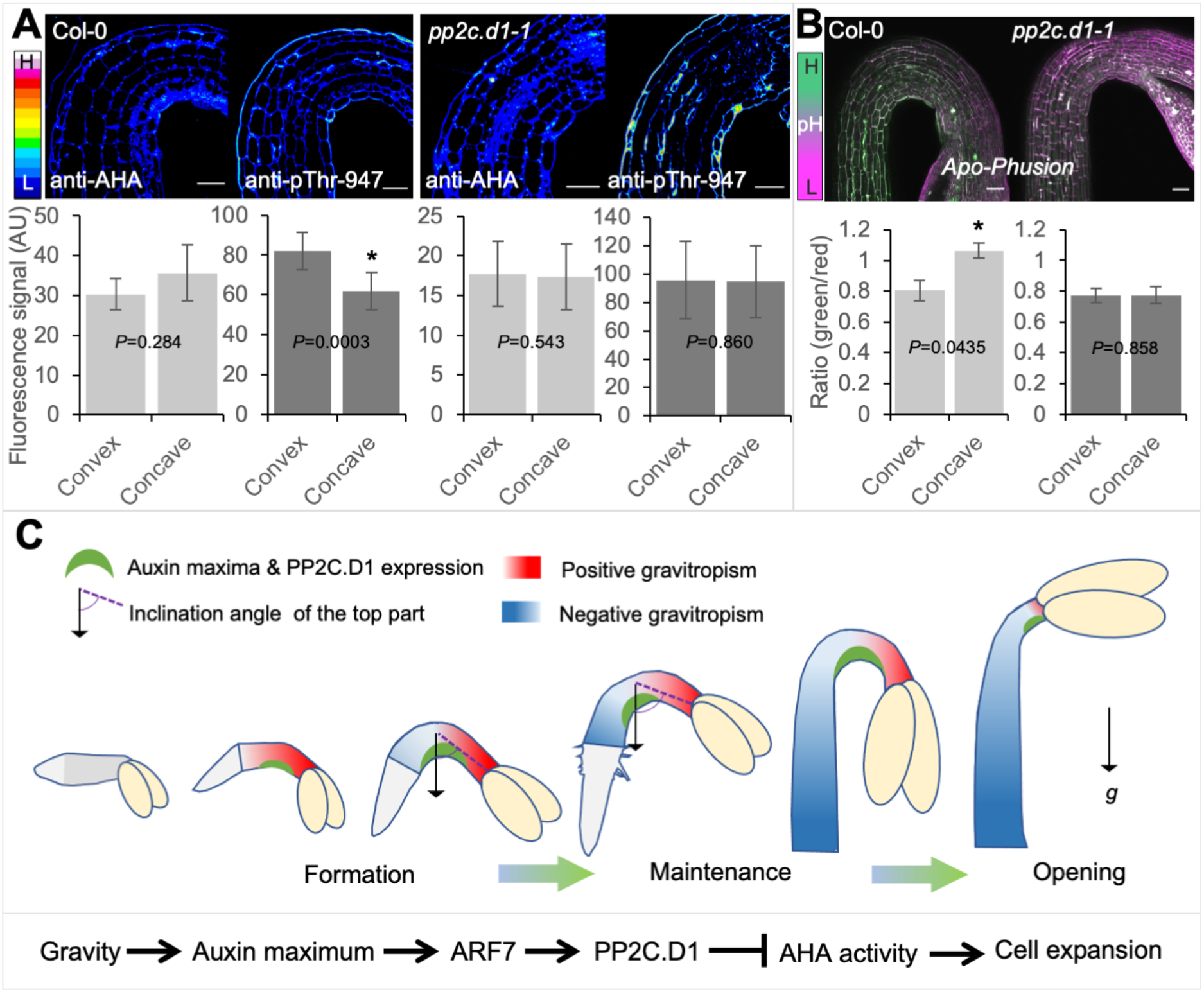
PP2C.D1 is required for asymmetric acid growth during apical hook development. **(A)** Immunolabeling and signal quantification of PM H^+^-ATPase and Thr^947^-phosphorylated PM H^+^-ATPase in WT and *pp2c*.*d1-1* mutant. Scale bars=50 µm. **(B)** Apoplast pH and signal quantification at the hook region using the *Apo-pHusion* marker line in WT and *pp2c*.*d1-1* backgrounds. Scale bars=50 µm. For **a** and **b**, details of the signal quantification can be found in the methods section. **(C)** Proposed model for hypocotyl elongation and apical hook development during etiolated seedling development. A molecular signaling pathway underlying auxin-mediated inhibition of cell elongation at the concave side of the hook is shown.

During plant gravitropic responses, shoots bend upwards (negative gravitropism) while roots bend downwards (positive gravitropism)(*39*). The basis of this opposite gravitropism is the regulation by auxin, which promotes cell elongation in shoots, while inhibiting it in roots. Since auxin also inhibits cell elongation in hypocotyls during early seedling development (Fig. 1 and fig. S2), hence, the hypocotyl can bend downwards in response to gravity. Consistent with this notion, hypocotyls bent upwards when the gravity-directed auxin flow was reversed in the *atlazy* mutants (Fig. 2, F and G). The downwards bending of the hypocotyl is the beginning of the hook formation.

Subsequently, as the auxin switches to promote cell elongation in the bottom cells (Fig. 1), the bottom part thus starts to bend upwards while the top part is still in a downward direction, and gradually causing the hook to be fully formed. Therefore, hook curvature is the consequence of the opposite gravitropic bending of the bottom and top parts of the young hypocotyl.

### Hook-expressed PP2C.D1 is specifically activated by auxin via ARF7

We next explored the molecular mechanism by which the gravity-induced auxin maxima at the concave side of the hook inhibits cell elongation. We paid particular attention to the PP2C.D1 protein phosphatase, because it is a negative regulator of cell elongation, exhibits a specific expression pattern in epidermal cells at the concave side of the hook(*16*) (Fig. 3A), and its expression is induced by IAA(*16*). Additionally, *pp2c*.*d1* mutants exhibit impaired apical hook development*(16, 40)* (Fig. 3, B and C) due to the release of growth inhibition at the concave side (fig. S5A). The highly specific expression of *PP2C*.*D1* coincides with the auxin maximum at the concave side of the hook (Fig. 3A). We found that the expression of *PP2C*.*D1::EGFP-GUS*(*16*) in the apical hook was auxin-regulated as shown in auxin-related mutants *pin3-3*(*27*), *arf7-1*(*41*) and *yuc1-D*(*26*) (fig. S5B). Furthermore, treatments with 10 µM IAA or 1 µM auxin efflux inhibitor N-(1-naphtyl) phtalamic acid (NPA), abolished the asymmetric expression of *PP2C*.*D1* (fig. S5C), indicating that asymmetric auxin distribution determines the pattern of *PP2C*.*D1* expression in the apical hook. Using RT-qPCR and western blot analyses, we further confirmed that *PP2C*.*D1* expression was induced by auxin in a dose-dependent manner (fig. S5, D and E).

The TMK1-mediated non-canonical auxin pathway was recently reported to regulate apical hook development through transcriptional regulation(*9*). We asked whether *PP2C*.*D1* expression is induced through the canonical TIR1/AFB auxin pathway(*25*) or non-canonical TMK1-mediated signaling. *PP2C*.*D1* expression in the hook region was reduced upon inhibiting the SCF^TIR1^ auxin receptor with auxinole (fig. S5F). In contrast, RT-qPCR analysis showed that auxin-induced *PP2C*.*D1* expression was comparable in WT and the *tmk1* mutant (Fig. 3D). These results suggest that *PP2C*.*D1* expression is regulated by the canonical TIR1/AFB pathway, rather than TMK1mediated signaling. ARF7 is an important transcriptional activator downstream of the TIR1/AFB receptors and is critical for apical hook development(*41*). The dramatically decreased *PP2C*.*D1* expression in the *arf7-1 arf19-1* mutant(*42*) (Fig. 3D) led us to reason that *PP2C*.*D1* may be directly regulated by ARF7. We performed chromatin immunoprecipitation (ChIP) experiments followed by qPCR using *arf7 arf19 pARF7::ARF7-GFP* seedlings(*43*). Strong ARF7-GFP enrichment was detected within a *PP2C*.*D1* promoter region (P4) containing two inverted auxin response elements (AuxRE)(*44*) (Fig. 3E). Together, these observations indicate that auxin induces *PP2C*.*D1* expression at the concave side of the hook through ARF7.

### PP2C.D1 controls asymmetric acid growth for apical hook development

*PP2C*.*D1* overexpression inhibits the activity of PM H^+^-ATPases(*15*) by dephosphorylating the penultimate threonine residue (Thr-947 in AHA2)(*14*). However, the lack of hypocotyl elongation defects in *pp2c*.*d1* mutants^20^ indicates that PP2C.D1 doesn’t play an important role in hypocotyl elongation. Rather, the specific *PP2C*.*D1* expression at the concave side of the hook suggests that PP2C.D1 predominantly inhibits PM H^+^-ATPase activity in these cells, thereby causing differential efflux of protons and altered apoplastic pH across the hook. We analyzed the *in vivo* distribution pattern of PM H^+^-ATPase at the hook region by immunolabeling with H^+^-ATPase and pThr-947 antibodies(*45*). In wild-type, while PM H^+^-ATPase was nearly equally distributed across the hook epidermis, we observed a higher accumulation of phosphorylated H^+^-ATPase at the convex side (Fig. 4A and fig. S6A). Consistently, using the genetically encoded apoplastic pH sensor *Apo-pHusion*(*46*), the pH at the convex side was lower than that at the concave side (Fig. 4B), suggesting that differential activation of proton pumps and the consequent asymmetric acid growth between the concave and convex sides drives hook curvature. Consistent with this possibility and the fact that SAUR19 promotes H^+^-ATPase activation(*15*), *35S:Strep-SAUR19* expression abolished the asymmetric distribution of phosphorylated proton pumps (fig. S6B), and conferred severe defects in hook development (Fig. 3, B and C). The hook defects in these seedlings correlated with increased epidermal cell length at both sides of the hook (fig. S6, C and D). Similar defects were also seen in the *open stomata2* (*ost2-2*) mutant (Fig. 3, B and C; fig. S6, C and D), which contains a constitutively active allele of the PM H^+^-ATPase encoded by AHA1(*47*). Importantly, asymmetries in PM H^+^-ATPase phosphorylation and apoplastic pH were also abolished in the *pp2c*.*d1-1* mutant (Fig. 4, A and B). These results demonstrate that differential acid-growth across the hook is established through PP2C.D1-mediated inhibition of PM H^+^ATPase activity at the concave side.

Together, these results indicate that PP2C.D1 acts downstream of TIR1/AFB-based canonical auxin signaling to inhibit cell elongation at the concave side of the hook. At lower concentrations (e.g. in the elongating part of the hypocotyl), auxin induces *SAUR* expression and therefore inhibits the PP2C.D2/5/6 phosphatases to promote cell elongation(*15, 16*) (fig. S7). By contrast, high auxin concentrations (e.g. at the concave side of the hook) induce *PP2C*.*D1* expression allowing the system to bypass SAUR-regulation and inhibit cell elongation through PP2C.D1 dephosphorylation of PM H^+^-ATPase (fig. S7). Interestingly, a recent study found that SAUR17 has a high affinity for PP2C.D1 without inhibiting its activity(*48*). This allows SAUR17 to shield PP2C.D1 from inhibition caused by other SAUR proteins, and could serve to reinforce the PP2C.D1-mediated H^+^-ATPase inhibition resulting from localized phosphatase expression.

## Conclusions

Our findings suggest a mechanistic framework for auxin-regulated hypocotyl elongation and apical hook development during seedling emergence (Fig. 4C). During early etiolated seedling development, high levels of auxin restrain cell elongation in the hypocotyl. This delay in growth coincides with the beginning of apical hook formation, as simultaneously, gravity generates an auxin maximum on one side of the hypocotyl which induces high *PP2C*.*D1* expression. Consequently, growth inhibition is strengthened at the lower side through PP2C.D1-dependent inhibition of PM H^+^-ATPase activity, and released at the opposite side, causing the hypocotyl to bend toward the gravity vector and form the hook. As the bottom cells of the hypocotyl slowly elongate and cell volumes increase, auxin levels fall below the threshold for inhibition and begin to promote hypocotyl elongation against the gravity vector. This in turn increases the inclination angle of the top of the hypocotyl to the direction of gravity (Fig. 4C). With the bottom part of the hypocotyl having a more vertical orientation, the gravity-induced auxin maximum is shifted upwards, moving the hook on its way. This is supported by the observations that the PP2C.D1GFP signal moved upwards (Fig. 3A) and the number of cells above the hook tip dropped during hook development (fig. S8). As the hypocotyl cells elongate acropetally, the proportion of the lower part of the hypocotyl that is negatively responding to gravity increases whilst the percentage of the upper part that is bending to the gravity vector decreases (fig. S8). Finally, when the acropetal wave of hypocotyl elongation reaches the very top cells, the hook starts to open.

This study provides critical insights into understanding how hypocotyl elongation and apical hook development are coordinated for successful seedling emergence. Auxin’s biphasic effect on cell elongation underlies this process. Early inhibition of hypocotyl elongation may facilitate emergence, as elongation prior to hook formation would increase kinetic friction, hindering soil penetration and potentially increasing mechanical damage to the cotyledons and shoot meristem. Consistent with this notion, *SAUR19* overexpression seedlings and *pp2c*.*d1* mutants displayed substantial reductions in soil emergence compared to WT controls (fig. S9). Coordination of elongation and hook development enables the seedling to, safely and efficiently, emerge from the soil and initiate phototropic development.

## Acknowledgments

We thank Edgar Spalding for providing the seeds of *atlazy* mutants, Hidehiro Fukaki and Masahiko Furutani for providing the *arf7 arf19 pARF7::ARF7-GFP* seeds, Toshinori Kinoshita for providing α-AHA and α-pThr-947 antisera, Ken-ichiro Hayashi for providing auxinole, and Clay Carter for providing his real-time PCR system for our qPCR experiments. We also thank the College of Biological Sciences Imaging Center for assistance with confocal microscopy.

## Author contributions

Conceptualization: MD, FB, WMG

Methodology: MD, FB, AS, MT

Investigation: MD, FB, AS, MT, YL, XZ, JHW, HR

Visualization: MD, FB

Funding acquisition: WMG, JDC

Project administration: WMG, MD

Supervision: WMG

Writing – original draft: MD

Writing – review & editing: MD, WMG, FB

M.D., F.B., and W.M.G. designed the research strategy. M.D., F.B., Y.L., X.Z., J.H.W. and H.R. performed experiments. A.S. and M.T. performed the IAA quantification. J.C. assisted with the IAA quantification. C.L. assisted with ChIP experiments and hosted M.D. to perform collaborative study in his laboratory. M.D., F.B., J.D.C., and W.M.G. analysed the data. M.D., W.M.G. and F.B. wrote the paper.

## Data and materials availability

The main data supporting the findings of this study are available within the paper and its supplementary materials. Additional data are available from the corresponding authors upon request. Correspondence and requests for materials should be addressed to C.L. or W.M.G.

## Supplementary Materials

Materials and Methods

Figs. S1 to S9

Table S1

References (*49*–*56*)

Movies S1 to S4

## References and Notes

1. E. Gendreau et al., Cellular basis of hypocotyl growth in Arabidopsis thaliana. Plant Physiology 114, 295–305 (1997).

2. C. Darwin, F. Darwin, The power of movement in plants. New York (Republished 1892), (1881).

3. V. Raz, J. R. Ecker, Regulation of differential growth in the apical hook of Arabidopsis. Development 126, 3661–3668 (1999).

4. M. Du, E. P. Spalding, W. M. Gray, Rapid auxin-mediated cell expansion. Annual Review of Plant Biology 71, 379–402 (2020).

5. H. Li, P. Johnson, A. Stepanova, J. M. Alonso, J. R. Ecker, Convergence of signaling pathways in the control of differential cell growth in Arabidopsis. Developmental Cell 7, 193–204 (2004).

6. C. Beziat, J. Kleine-Vehn, The road to auxin-dependent growth repression and promotion in apical hooks. Current Biology 28, R519–r525 (2018).

7. F. Vandenbussche et al., The auxin influx carriers AUX1 and LAX3 are involved in auxin-ethylene interactions during apical hook development in Arabidopsis thaliana seedlings. Development 137, 597–606 (2010).

8. P. Zadnikova et al., Role of PIN-mediated auxin efflux in apical hook development of Arabidopsis thaliana. Development 137, 607–617 (2010).

9. M. Cao et al., TMK1-mediated auxin signalling regulates differential growth of the apical hook. Nature 568, 240–243 (2019).

10. K. V. Thimann, Auxins and the inhibition of plant growth. Biological Reviews 14, 314–337 (1939).

11. Q. Zhu et al., Root gravity response module guides differential growth determining both root bending and apical hook formation in Arabidopsis. Development 146, (2019).

12. Y. Wang, H. Guo, On hormonal regulation of the dynamic apical hook development. New Phytologist 222, 1230–1234 (2019).

13. D. L. Rayle, R. E. Cleland, The acid growth theory of auxin-induced cell elongation is alive and well. Plant Physiology 99, 1271–1274 (1992).

14. K. Takahashi, K. Hayashi, T. Kinoshita, Auxin activates the plasma membrane H^+^-ATPase by phosphorylation during hypocotyl elongation in Arabidopsis. Plant Physiology 159, 632–641 (2012).

15. A. K. Spartz et al., SAUR inhibition of PP2C-D phosphatases activates plasma membrane H^+^-ATPases to promote cell expansion in Arabidopsis. The Plant Cell 26, 2129–2142 (2014).

16. H. Ren, M. Y. Park, A. K. Spartz, J. H. Wong, W. M. Gray, A subset of plasma membrane-localized PP2C.D phosphatases negatively regulate SAUR-mediated cell expansion in Arabidopsis. PLoS Genetics 14, e1007455 (2018).

17. M. Fendrych et al., Rapid and reversible root growth inhibition by TIR1 auxin signalling. Nature Plants 4, 453–459 (2018).

18. E. Barbez, K. Dünser, A. Gaidora, T. Lendl, W. Busch, Auxin steers root cell expansion via apoplastic pH regulation in Arabidopsis thaliana. Proceedings of the National Academy of Sciences 114, E4884–E4893 (2017).

19. F. Bou Daher et al., Anisotropic growth is achieved through the additive mechanical effect of material anisotropy and elastic asymmetry. eLife 7, (2018).

20. C.-Y. Liao et al., Reporters for sensitive and quantitative measurement of auxin response. Nature Methods 12, 207–210 (2015).

21. M. Narukawa-Nara et al., Aminooxy-naphthylpropionic acid and its derivatives are inhibitors of auxin biosynthesis targeting l-tryptophan aminotransferase: structure-activity relationships. The Plant Journal 87, 245–257 (2016).

22. K. Hayashi et al., Rational design of an auxin antagonist of the SCF^TIR1^ auxin receptor complex. ACS Chemical Biology 7, 590–598 (2012).

23. A. N. Stepanova et al., TAA1-mediated auxin biosynthesis is essential for hormone crosstalk and plant development. Cell 133, 177–191 (2008).

24. Y. Tao et al., Rapid synthesis of auxin via a new tryptophan-dependent pathway is required for shade avoidance in plants. Cell 133, 164–176 (2008).

25. N. Dharmasiri et al., Plant development is regulated by a family of auxin receptor F box proteins. Developmental Cell 9, 109–119 (2005).

26. Y. Zhao et al., A role for flavin monooxygenase-like enzymes in auxin biosynthesis. Science 291, 306–309 (2001).

27. J. Friml, J. Wisniewska, E. Benkova, K. Mendgen, K. Palme, Lateral relocation of auxin efflux regulator PIN3 mediates tropism in Arabidopsis. Nature 415, 806–809 (2002).

28. I. R. MacDonald, D. C. Gordon, J. W. Hart, E. P. Maher, The positive hook: the role of gravity in the formation and opening of the apical hook. Planta 158, 76–81 (1983).

29. A. B. Myers, R. D. Firn, J. Digby, Gravitropic sign reversal-a fundamental feature of the gravitropic perception or response mechanisms in some plant organs. Journal of Experimental Botany 45, 77–83 (1994).

30. F. Migliaccio, O. Micciulla, S. Ferrari, Hook formation in sunflower seedlings is directed by both positive gravitropism and a form of circumnutation. Plant Biosystems 132, 11–16 (1998).

31. H. Chauvet, O. Pouliquen, Y. Forterre, V. Legué, B. Moulia, Inclination not force is sensed by plants during shoot gravitropism. Scientific Reports 6, 35431 (2016).

32. O. Pouliquen et al., A new scenario for gravity detection in plants: the position sensor hypothesis. Physical Biology 14, 035005 (2017).

33. M. G. Heisler et al., Patterns of auxin transport and gene expression during primordium development revealed by live imaging of the Arabidopsis inflorescence meristem. Current Biology 15, 1899–1911 (2005).

34. H. Fukaki et al., Genetic evidence that the endodermis is essential for shoot gravitropism in Arabidopsis thaliana. The Plant Journal 14, 425–430 (1998).

35. M. Nakamura, T. Nishimura, M. T. Morita, Bridging the gap between amyloplasts and directional auxin transport in plant gravitropism. Current Opinion in Plant Biology 52, 54–60 (2019).

36. T. Yoshihara, E. P. Spalding, LAZY genes mediate the effects of gravity on auxin gradients and plant architecture. Plant Physiology 175, 959–969 (2017).

37. J. Friml et al., Efflux-dependent auxin gradients establish the apical-basal axis of Arabidopsis. Nature 426, 147–153 (2003).

38. H. Rakusová et al., Polarization of PIN3-dependent auxin transport for hypocotyl gravitropic response in Arabidopsis thaliana. The Plant Journal 67, 817–826 (2011).

39. P. H. Masson et al., Arabidopsis thaliana: A Model for the Study of Root and Shoot Gravitropism. The Arabidopsis Book 1, e0043–e0043 (2002).

40. M. Sentandreu et al., Functional profiling identifies genes involved in organ-specific branches of the PIF3 regulatory network in Arabidopsis. The Plant Cell 23, 3974–3991 (2011).

41. R. M. Harper et al., The NPH4 locus encodes the auxin response factor ARF7, a conditional regulator of differential growth in aerial Arabidopsis tissue. The Plant Cell 12, 757–770 (2000).

42. Y. Okushima et al., Functional genomic analysis of the AUXIN RESPONSE FACTOR gene family members in Arabidopsis thaliana: Unique and overlapping functions of ARF7 and ARF19. The Plant Cell 17, 444–463 (2005).

43. J. Ito et al., Auxin-dependent compositional change in Mediator in ARF7-and ARF19-mediated transcription. Proceedings of the National Academy of Sciences 113, 6562–6567 (2016).

44. A. Freire-Rios et al., Architecture of DNA elements mediating ARF transcription factor binding and auxin-responsive gene expression in Arabidopsis. Proceedings of the National Academy of Sciences 117, 24557–24566 (2020).

45. Y. Hayashi et al., Biochemical characterization of in vitro phosphorylation and dephosphorylation of the plasma membrane H^+^-ATPase. Plant and Cell Physiology 51, 1186–1196 (2010).

46. K. S. Gjetting, C. K. Ytting, A. Schulz, A. T. Fuglsang, Live imaging of intra-and extracellular pH in plants using pHusion, a novel genetically encoded biosensor. Journal of Experimental Botany 63, 3207–3218 (2012).

47. S. Merlot et al., Constitutive activation of a plasma membrane H^+^-ATPase prevents abscisic acid-mediated stomatal closure. The EMBO Journal 26, 3216–3226 (2007).

48. J. Wang et al., SAUR17 and SAUR50 differentially regulate PP2C-D1 during apical hook development and cotyledon opening in Arabidopsis. The Plant Cell 32, 3792–3811 (2020).

49. A. K. Spartz et al., The SAUR19 subfamily of SMALL AUXIN UP RNA genes promote cell expansion. The Plant Journal 70, 978–990 (2012).

50. L. Willis et al., Cell size and growth regulation in the Arabidopsis thaliana apical stem cell niche. Proceedings of the National Academy of Sciences 113, E8238–E8246 (2016).

51. Q. Tang, P. Yu, M. Tillmann, J. D. Cohen, J. P. Slovin, Indole-3-acetylaspartate and indole-3-acetylglutamate, the IAA-amide conjugates in the diploid strawberry achene, are hydrolyzed in growing seedlings. Planta 249, 1073–1085 (2019).

52. X. Liu, A. D. Hegeman, G. Gardner, J. D. Cohen, Protocol: High-throughput and quantitative assays of auxin and auxin precursors from minute tissue samples. Plant Methods 8, 31 (2012).

53. J. D. Cohen, B. G. Baldi, J. P. Slovin, C(6)-[benzene ring]-indole-3-acetic acid: A new internal standard for quantitative mass spectral analysis of indole-3-acetic acid in plants. Plant Physiology 80, 14–19 (1986).

54. L. S. Barkawi, Y. Y. Tam, J. A. Tillman, J. Normanly, J. D. Cohen, A high-throughput method for the quantitative analysis of auxins. Nature Protocols 5, 1609–1618 (2010).

55. M. Du et al., MYC2 orchestrates a hierarchical transcriptional cascade that regulates jasmonate-mediated plant immunity in tomato. The Plant Cell 29, 1883–1906 (2017).

56. M. Postma, J. Goedhart, PlotsOfData-A web app for visualizing data together with their summaries. PLOS Biology 17, e3000202 (2019).

