## Supplemental Materials for "Biphasic Control of Cell Expansion by Auxin Coordinates Etiolated Seedling Development"

**This PDF file includes:**

Materials and Methods
Figs. S1 to S9
Tables S1
Captions for Movies S1 to S4

**Materials and Methods**

### Plant materials

All Arabidopsis (*Arabidopsis thaliana*) lines used in this study were in the Col-0 background except the *R2D2* reporter(*20*), which is in the Columbia-Utrecht (Col-utr) ecotype. The transgenic and mutant lines used have been described previously: *35S::StrepII-SAUR19(49)*, *PP2C.D1::PP2C.D1-GFP(16)*, *PP2C.D1::EGFP-GUS(16)*, *DR5::VENUS-NLS(33)*, *MYR-YFP(50)*, *PIN3::PIN3-GFP(8)*, *Apo-pHusion(46)*, *tmk1(9)*, *yuc1-D(26)*, *wei8-1 tar2-1(23) (24)*, *tir1 afb2(25)*, *atlazy2,3,4(36)*, *atlazy1,2,3,4(36)*, *scr-3(34)*, *shr-2(34)*, *pp2c.d1-1(40)*, *pp2c.d1-2(40)*, *ost2-2(47)*, *pin3-3(27)*, *arf7-1(41)*, *arf7-1 arf19-1(42)*, *arf7 arf19 ARF7::ARF7-GFP(43)*.

### Growth conditions

All Arabidopsis materials were grown in continuous darkness at 22 °C, unless otherwise indicated. For ChIP-qPCR analysis, seeds were sterilized with 30% bleach for 15 minutes, washed 3 times with sterile water, and sown on ½ Murashige & Skoog (MS, PhytoTech Labs, M524) plates containing 1% sucrose (Macron, 8360-06) and 0.6% agargel (Sigma-Aldrich, A3301). Media pH was adjusted to 5.7 using KOH (Fisher Chemical, P250-1). After sowing, the seeds were left at 4°C for 48 hours in darkness, then transferred to white light for 6 hours to stimulate germination, and subsequently to darkness and left at 22°C for the desired time. For other experiments, seeds were sterilized with 70% ethanol for 5 minutes and germinated on ½ MS plates containing 0.6% agargel (pH 5.7). Germination was defined as the time when the radicle broke through the endosperm (0HPG). At this time, seedlings were selected and aligned horizontally or vertically on ½ MS plates containing 1% sucrose and 0.6% agargel (pH 5.7). The plates were wrapped with two layers of foil to simulate constant darkness.

### Quantification of angles of hook curvature

The angle of hook curvature is defined as 180° minus the angle between the tangential of the apical part with the axis of the lower part of the hypocotyl(*8*). For quantifying the hook angles of *scr-3*, *shr-2*, *atlazy2,3,4* and *atlazy1,2,3,4* mutants, germinated seeds were initially placed at a horizontal orientation on ½ MS plates containing 1% sucrose and 0.6% agargel (pH 5.7) at 0HPG, which were wrapped with two layers of foil and kept vertically in the growth chamber. Pictures of the seedlings were taken at 24HPG for measuring the hook angles using ImageJ (NIH, Bethesda, MD, USA). For measuring the hook angles in the rotation experiment, germinated seeds were initially placed horizontally. The wrapped plates were rotated 180 degrees once per hour (rotations were done at 1HPG, 2HPG, 3HPG…11HPG). Pictures of the seedlings were taken at 12HPG for measuring the hook angles. For analyzing the kinematics of apical hook development, germinated seeds were initially placed at a vertical orientation on ½ MS plates with 1% sucrose and 0.6% agargel (pH 5.7) at 0HPG, which were kept vertically in darkness at 22°C. Development of seedlings was recorded at 15 min intervals with an infrared light source (880 nm LED; Advanced Illumination) by a Stingray F146B CCD camera (Allied Vision Technologies) equipped with a 18-108 mm macro video zoom lens with an infrared long pass filter. Images for selected time points were extracted and angles of hook curvature were measured using ImageJ. Frames with six-hour intervals were imported in ImageJ and color coded using the (Rainbow RGB-LUT projection) in the temporal-color coding tool.

### Measurement of epidermal cell length

For measuring the cell length of hypocotyl epidermal cells during etiolated seedlings development, Arabidopsis hypocotyls were synchronized by selecting seeds at germination (0HPG; radical emergence from endosperm). Germinated seeds were placed at a vertical orientation and then used for measurements at different timepoints. Seedlings were stained with 0.1mg/mL of propidium iodide (PI, Molecular Probes, P3566) for 10 minutes prior to imaging The outline of epidermal cells was imaged using a Leica DM5000B fluorescence microscope equipped with a 490-510nm excitation and 520-550nm band pass emission filter cube and a 20X air objective lens. Epidermal cell length was measured using ImageJ software. Analyses were performed on middle epidermal files with no cell division.

For measuring the epidermal cell length at the concave and convex side of the hook, germinated seeds were initially placed at a horizontal orientation at 0HPG. At 36HPG, hooks were dissected using custom made blades, placed in longitudinal slits on 1.5% agar pads and imaged using transmitted light on a Leica DM5000B microscope. Measurements were done using ImageJ.

### Immunolabelling

Seedlings were fixed in a PBS solution containing 2% formaldehyde (Acros 11969) and 2.5% glutaraldehyde (Sigma-Aldrich, G6257) under vacuum for one hour and left at 4°C overnight. Samples were washed twice in PBS then dehydrated in a series of increasing ethanol concentrations. Samples were then embedded in medium grade LR White (London Resin Company) which was left to polymerize at 60°C overnight. 0.5µm thin sections were generated using glass knives on a Leica UltraCut microtome. Sections were blocked in PBS solution containing 2% bovine serum albumin (BSA) (Sigma-Aldrich, 2153) for one hour. The primary antibodies (Anti-AHA and anti-pThr947) were diluted 500 times in PBS-BSA and left on the samples overnight at 4°C. Samples were then washed four times for five minutes each in PBS-BSA. Samples were then incubated for three hours in DyLight 488 donkey-anti-rabbit secondary antibody (Biolegend, 406404) which was diluted 400 times in PBS-BSA. Samples were then washed six times for 5 minutes each in PBS. Sections were mounted in citifluor CFM-3 (Electron Microscopy Sciences) and slides were sealed with nail polish and imaged within two days from mounting them.

### Confocal imaging

For *R2D2* imaging, germinated seeds were placed at a vertical orientation at 0HPG and the seedlings were examined at indicated times. For IAA treatment, 12HPG old seedlings were incubated with DMSO (Mock) or 100 nM IAA for 30 minutes before imaging. For KOK2153 treatment, germinated seeds were grown on the plates supplemented with DMSO (Mock) or 10 μM KOK2153 (KOK; custom synthesis by LabSeeker, Wujiang City, China) for 12 hours before imaging. For *PIN3::PIN3-GFP* imaging, germinated seeds were placed at a horizontal orientation at 0HPG and the seedlings were examined at 6HPG. For *PIN3::PIN3-GFP* sample preparation, seedlings were dissected out of the seed coat and cut perpendicular to the hypocotyl axis using custom designed pins and blades. They were then placed in an agar pad with the cut side facing the outside. A drop of water was applied and the samples were covered with a coverslip and imaged using a confocal microscope. For *Apo-pHusion* imaging, germinated seeds were placed at a horizontal orientation at 0HPG and the seedlings were examined at 36HPG. For *MYR-YFP* imaging, germinated seeds were placed at a horizontal orientation at 0HPG and the seedlings were examined at different timepoints. The tip of the hook was determined using the kappa plugin in FIJI. Briefly, the points with highest curvature on the concave and convex sides were determined and a line joining them indicates the middle of the hook.

Confocal imaging was conducted using a Nikon A1si confocal microscope. For *DR5::GFP*, *PIN3::PIN3-GFP* and DyLight-488 imaging, a 20X, 0.75 numerical aperture objective was used. Samples were excited with a 480nm laser and the emitted laser was collected between 500 and

550nm. For *MYR-YFP* and *DR5::VENUS-NLS*, a 10X, 0.45 numerical aperture objective was used. A 514.5nm excitation laser and a 510-570mm emission filter were used. For *R2D2* and *Apo-pHusion*, images were acquired using a 10X, 0.45 numerical aperture objective. A 488nm excitation laser and 500-550mn emission were used for the first channel and 562 excitation and 570-620nm emission filter were used for the second channel.

### Image processing and analysis

All confocal images were processed using ImageJ. Maximum projection images were generated, and background noise was subtracted using a 50-pixel rolling ball radius. For immunolabelling images, a 16-color look up table was applied. A 3-pixel wide line was used in ImageJ to plot the fluorescence intensity profile on the outer periclinal walls of the epidermal cells on the concave and the convex sides. For processing *R2D2* images, after background removal, a Gaussian blur was applied with a 2.0 sigma (radius). The red channel was then divided by the yellow channel using the image calculator function to generate an image that was used to quantify the signal ratio in the nucleus. The signal was measured using the average intensity per nucleus and data were exported to excel. For the *Apo-pHusion* images, a 20-pixel wide line was used in ImageJ to plot the fluorescence intensity profile of the green and red channels on the epidermal cells both at the concave and the convex sides of the hook. For *PIN3::PIN3-GFP*, a 16-color look up table was applied. A 3-pixel wide line was used in ImageJ to plot the fluorescence intensity profile on the outer periclinal walls of the endodermis. All data were then exported to and analyzed in Microsoft Excel.

### Quantification of endogenous IAA

To quantify the IAA levels in the elongating hypocotyls at different times, seedlings were grown on ½ MS plates containing 1% sucrose and 1.5% agargel (pH 5.7). The shoot meristem, cotyledons and root were removed before harvesting the hypocotyls for analysis. For measuring the IAA levels in the bottom and top part, the hypocotyls at 36 HPG were cut into two halves and analyzed separately.

Free IAA was measured by isotope dilution essentially as previously described(*51*). The samples were analyzed by LC–MS/MS at high resolution. [$^{13}C_6$]IAA internal standard was added with 2-propanol/buffer(*52*) to plant tissue samples weighing 7-21 mg, which were then homogenized and incubated for 1 h on ice. Samples were then diluted with water, centrifuged, and IAA was extracted from the supernatant using amino and polymethylmethacrylate epoxide (PMME) solid phase extraction resins in Top Tips spin tips (Glygen, Columbia, MD, USA). After elution from PMME tips with methanol, sample volumes were reduced to approximately 20 μL and transferred to autosampler vials for LC–MS/MS analysis with a Dionex Ultimate 3000 RSLC HPLC coupled to a hybrid quadrupole Orbitrap Q Exactive mass spectrometer (Thermo Scientific). 8-10 μL of extract was injected onto a 50 × 2.1 mm Force C18 column with 1.8 μm particle size (Restek, Bellefonte, PA, USA) and run with a solvent gradient of 0.1% formic acid in water (solvent A) and 0.1% formic acid in acetonitrile (solvent B) at a flow rate of 0.4 mL $min^{-1}$. Gradient parameters were as follows: − 1 to 0 min, 5% B; 0–3 min, 5–20% B; 3–6 min, 20–80% B; and 6–6.5 min, 80% B. Mass spectrometry data were collected in the parallel reaction monitoring (PRM) scan mode with the [M + 1] for IAA and [$^{13}C_6$]IAA at *m/z* 176.08 and 182.1, respectively, in the inclusion list. Extracted ion chromatogram peaks for 130.0641-130.0661 *m/z* (corresponding to unlabeled quinolinium ion) and 136.0843-136.0863 *m/z* ([$^{13}C_6$] quinolinium produced from [$^{13}C_6$]IAA internal standard) were selected at 4.4-4.8 min. retention time, and peak areas were used to calculate endogenous IAA levels by isotope dilution(*53, 54*).

For calculating the IAA concentration, the IAA amount was divided by the fresh weight or the total volume of the hypocotyls, which was estimated by hypocotyl number times hypocotyl volume. We considered the hypocotyl as a truncate cone, whose volume was calculated using the formula: volume = $(1/3) * \pi * L * ((D/2)^2 + (d/2)^2 + (D/2) * (d/2))$ , where L is the hypocotyl length, D is the diameter of the base of the hypocotyl, and d is the diameter of the top of the hypocotyl.

### GUS staining

*SAUR22::GUS and PP2C.D1::EGFP-GUS* seedlings at indicated times were used for GUS staining. Seedlings were incubated in the staining buffer containing 0.1 M sodium phosphate buffer (pH 7), 1 mM $K_4Fe(CN)_6$, 1 mM $K_3Fe(CN)_6$, 0.1% Triton X-100 and 1 mg/ml X-Gluc for 2 hours at 37°C. GUS expression patterns were imaged with an Olympus SZX12 dissecting microscope using SPOT Advanced imaging software.

### Phytohormone and chemical treatment

Germinated seeds were grown on 1/2 MS plates containing 1% sucrose and 0.6% agargel (pH 5.7) supplemented with various chemicals: 100 nM IAA (Chem-Implex International, 00188) or 1 µM NPA (Duchefa Biochemie), 10 µM KOK2153 (also called Pyruvamine 2153)(*21*), or 10 µM auxinole. For gene expression, western blot and ChIP assays, we transferred 3-day-old dark-grown seedlings to 1/2 MS liquid medium containing 1% sucrose and different concentrations of IAA for 2 hours.

### RNA extraction and RT-qPCR analysis

Dark-grown seedlings were treated with different concentrations of IAA in 1/2 MS liquid medium for 2 hours. Total RNA was extracted from harvested seedlings using the NucleoSpin RNA Plant kit (Macherey-Nagel) and the quality of the total RNA was determined using an Implen NanoPhotometer P330. 2 μg of total RNA were used to synthesize the first strand cDNA with the M-MLV reverse transcriptase kit (Promega, M1701). RT-qPCR was performed on a StepOnePlus Real-Time PCR System (Applied Biosystems) with the Brilliant III Ultra-Fast SYBR Green QPCR Master Mix (Agilent Technologies, 600882). The expression levels of genes were normalized to that of the *ACTIN7* gene. Primer information is given in table S1.

### Protein extraction and Western Blot

Three-day-old dark-grown *PP2C.D1::PP2C.D1-GFP* seedlings were treated with different concentrations of IAA in 1/2 MS liquid medium containing 1% sucrose for 2 hours. Microsomal fractions were prepared by two-phase partitioning as previously described(*15*). Twenty micrograms of microsomal proteins were mixed with SDS-PAGE sample buffer, separated by SDS-PAGE, and blotted to nitrocellulose. The proteins were detected by western blot with anti-GFP primary antibody (Covance, MMS-118R). A non-specific band detected by the antibody was served as a loading control.

### ChIP-qPCR assay

Three-day-old dark-grown *arf7 arf19 ARF7::ARF7-GFP* seedlings treated with or without 10 μM IAA were used for ChIP-qPCR analysis. The ChIP-qPCR assay was conducted as described previously(*55*). The ChIP signal was quantified by qPCR. The primers used in qPCR were designed to amplify various regions of the *PP2C.D1* promoter. Each ChIP value was normalized to its respective input DNA value, and enrichment of DNA is shown as the percentage of input.

### Statistical analysis

All box plots were generated using the PlotsOfData(*56*) or ggplot2 in R studio. In the boxplots, the top, bottom and middle lines represent the 75th percentile, the 25th percentile and the median, respectively. Bar graphs and connecting lines were generated using Microsoft Excel. All the experiments were performed at least three times. The values were collected from three biological replicates. Student's t-test and Analysis of variance (ANOVA) were used for statistical analysis, using Microsoft Excel.

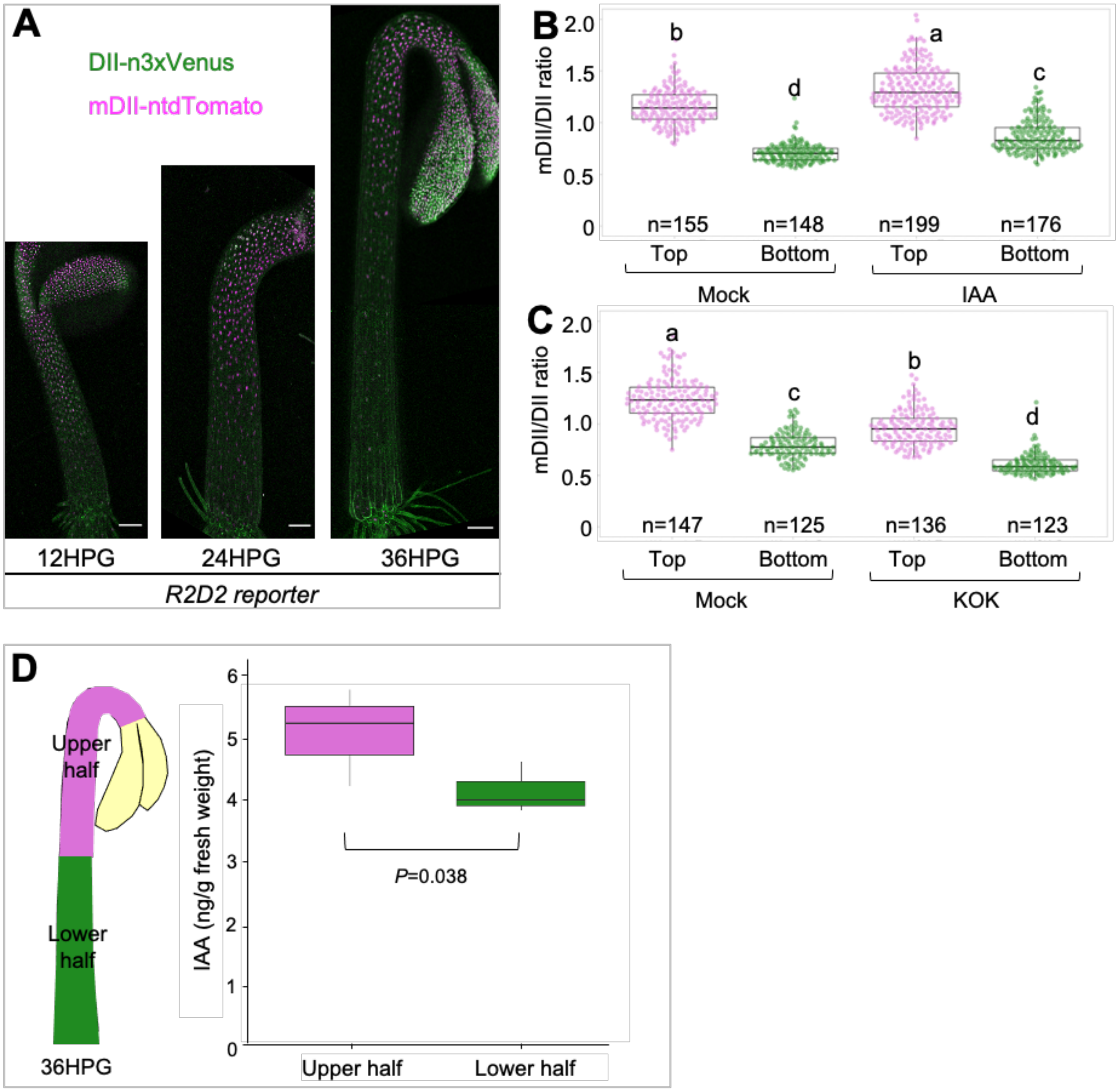


**Fig. S1. Spatial distribution of auxin signaling in the hypocotyl epidermis. (A)** n3×Venus (green) and ntdTomato (magenta) fluorescence signal overlays in hypocotyls of *R2D2* seedlings 12-36 hours post germination (HPG). **(B and C)** Quantification of the mDII/DII ratio at the top and bottom parts of *R2D2* hypocotyls treated with IAA or KOK2153. For IAA treatment (**B**), 12HPG seedlings were incubated with DMSO (Mock) or 100 nM IAA for 30 minutes prior to imaging. For KOK2153 treatment **(C)**, germinated seeds were grown on plates supplemented with DMSO (Mock) or KOK2153 (KOK) for 12 hours prior to imaging. Different letters indicate statistical differences with $P$ value < 0.001. **(D)** Quantification of IAA in the upper and lower halves of hypocotyls at 36HPG. A schematic diagram shows the upper and lower halves of the hypocotyl used. A paired t-test was used to analyze the difference between the two halves and $P$ value was indicated.

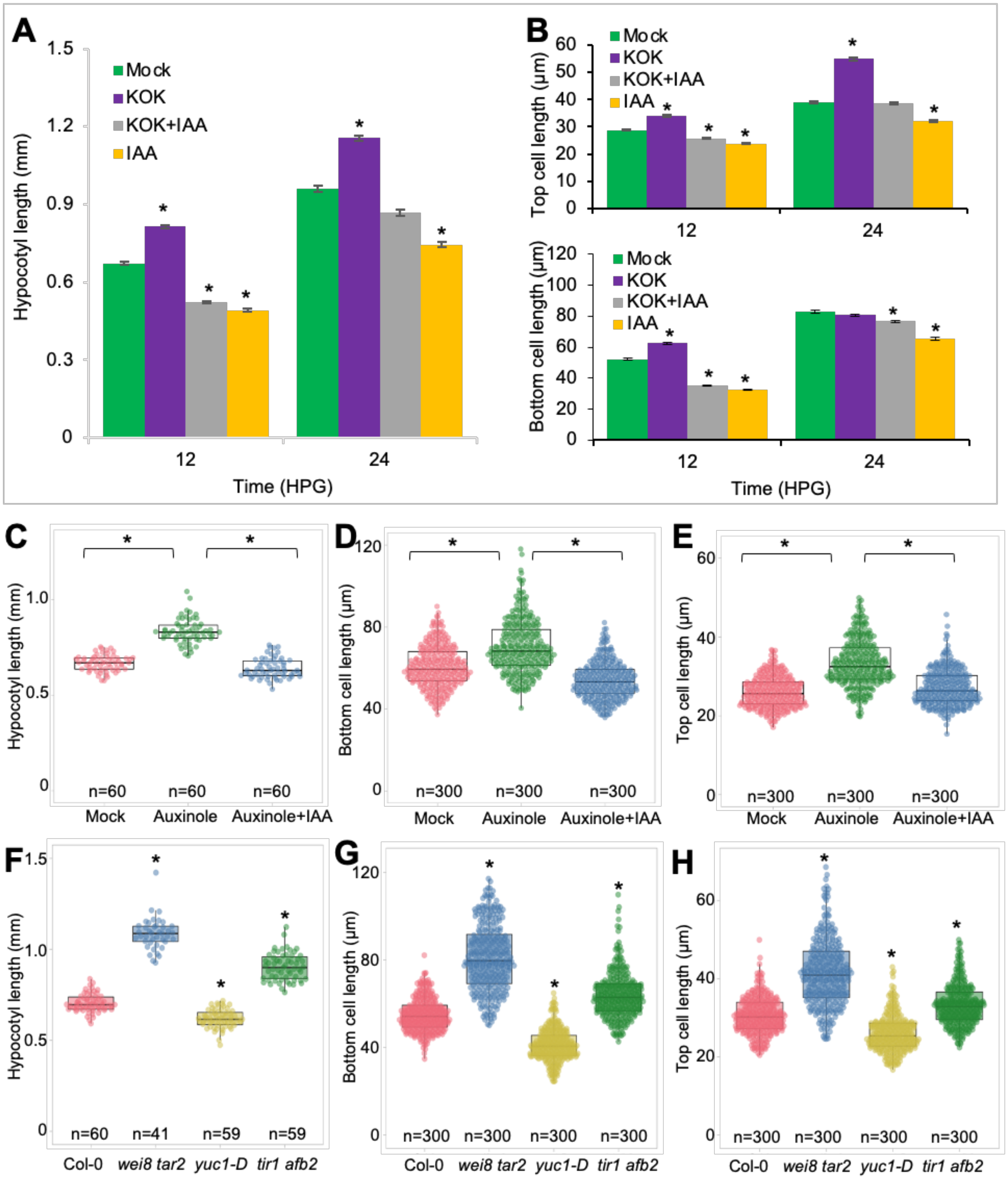


**Fig. S2. Auxin inhibits hypocotyl epidermal cell elongation during early etiolated development. (A)** Quantification of hypocotyl length during etiolated seedling development, n=60 hypocotyls. **(B)** Quantification of epidermal cell length of the bottom and top cells during etiolated seedling development, n=300 cells. For (**A** and **B**), germinated seeds were transferred to half-strength MS medium supplemented with DMSO (Mock), 10 μM KOK2153 (KOK), 100 nM IAA (IAA) or KOK2153 plus IAA (KOK+IAA). Hypocotyl and cell length were measured at the indicated time, values represent sample means ± s.e.m. from three replicates. * indicates *P* value < 0.01. **(C)** Quantification of hypocotyl length of etiolated seedlings grown on media with different treatments, n=60 hypocotyls. **(D** and **E)** Quantification of epidermal cell length of the bottom (**D**) and top (**E**) cells of etiolated hypocotyls grown on media with different treatments. For (**C** to **E)**, germinated seeds were transferred to half-strength MS media supplemented with DMSO (Mock), 10 μM auxinole (Auxinole) or auxinole plus IAA (Auxinole+IAA). Hypocotyl and cell length were measured at 12HPG, values represent sample means ± s.e.m. from three replicates. * indicates *P* value < 0.01. (**F** to **H)** Quantification of hypocotyl length (**F**) and epidermal cell length (**G** and **H**) of different genotypes. Germinated seeds of Col-0, *wei8 tar2*, *yuc1-D* and *tir1 abf2* were transferred to half-strength MS medium supplemented with 1% sucrose at 0HPG. Hypocotyl and cell lengths were measured at 12HPG, values represent sample means ± s.e.m. from three replicates. For (**B**, **D**, **E**, **G** and **H)**, five bottom cells (No. 2-6) and five top cells (No. 16-20) were used for quantification. * indicates *P* value < 0.01.

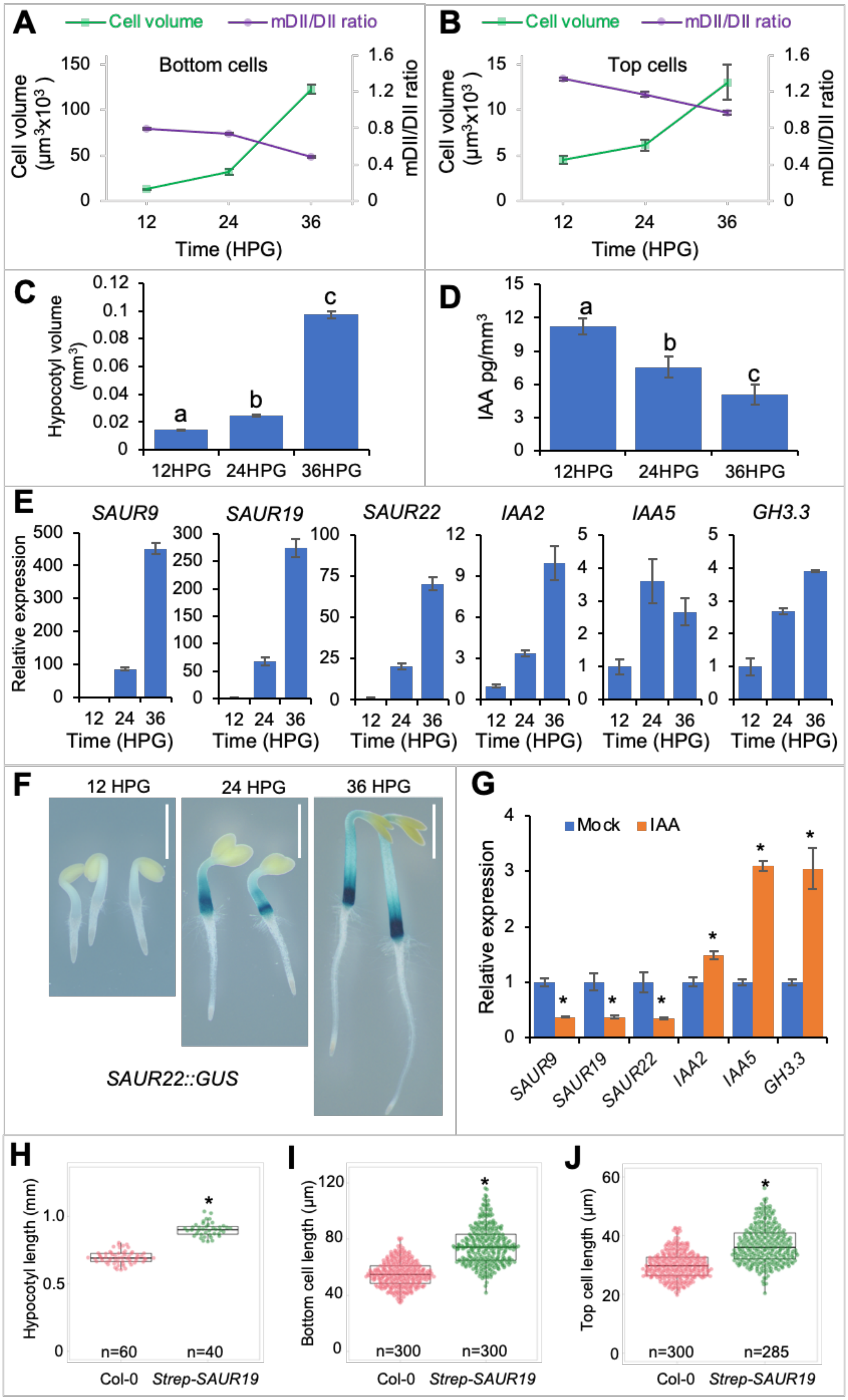
A
Cell volume
mDII/DII ratio
Bottom cells
Cell volume (μm³x10³)
mDII/DII ratio
Time (HPG)
B
Top cells
C
Hypocotyl volume (mm³)
12HPG
24HPG
36HPG
D
IAA pg/mm³
E
SAUR9
SAUR19
SAUR22
IAA2
IAA5
GH3.3
Relative expression
F
12 HPG
24 HPG
36 HPG
SAUR22::GUS
G
Mock
IAA
H
Hypocotyl length (mm)
n=60
n=40
Col-0
Strep-SAUR19
I
Bottom cell length (μm)
n=300
n=300
J
Top cell length (μm)
n=300
n=285

**Fig. S3. Decreased auxin levels correlate with increased cell volume and *SAUR* expression during hypocotyl elongation. (A and B)** decreased auxin levels correlate with increased cell volume of bottom (**B**) and top (**C**) cells of *R2D2* hypocotyls for the indicated time. Auxin levels were proxied by the inverse n3×Venus/ ntdTomato signal ratio (mDII/DII). Cells were assumed to have a cylindrical shape and cell volume were estimated by the formula for the volume of a cylinder [π*(width/2)^2*length]. Four cells (No. 2-5) from the bottom part and three cells (No. 18-20) from the top part were used for analysis. Values represent sample means ± s.e.m. from three replicates. **c**, Quantifications of hypocotyl volumes at indicated times. **(D)** Quantifications of endogenous IAA in the elongating hypocotyls at indicated times. Auxin concentrations were expressed in ng IAA per volume of hypocotyl ($mm^3$). Different letters indicate statistical differences between groups with $P$ value < 0.01 using ANOVA test. **(E)** Expression of auxin-responsive genes in etiolated hypocotyls at the indicated times. **(F)** *SAUR22::GUS* expression in etiolated hypocotyls at the indicated times. Scale bars=1 mm. **(G)** Expression of auxin-responsive genes in etiolated hypocotyls grown on different media. Germinated seeds were transferred to half-strength MS medium supplemented 1% sucrose and with DMSO (Mock) or 1 µM IAA (IAA) for growth. Expression analysis were performed at 36HPG. For **(E and G)**, expression levels of indicated genes were normalized against *ACTIN7* expression and presented as the fold change to the expression at 12HPG (**E**) or Mock (**G**). Data are means ± s.e.m. from three biological replicates. * indicates $P$ value < 0.01. **(H to J)** Quantification of hypocotyl length (**H**) and epidermal cell length (**I** and **J**) of Col-0 and *35S::Strep-SAUR19*. Germinated seeds were transferred to half-strength MS medium (with 1% sucrose) at 0HPG. Hypocotyl and cell lengths were measured at 12HPG, values represent sample means ± s.e.m. from three replicates. Five bottom cells (No. 2-6) and five top cells (No. 16-20) were used for quantification. * indicates $P$ value < 0.01.

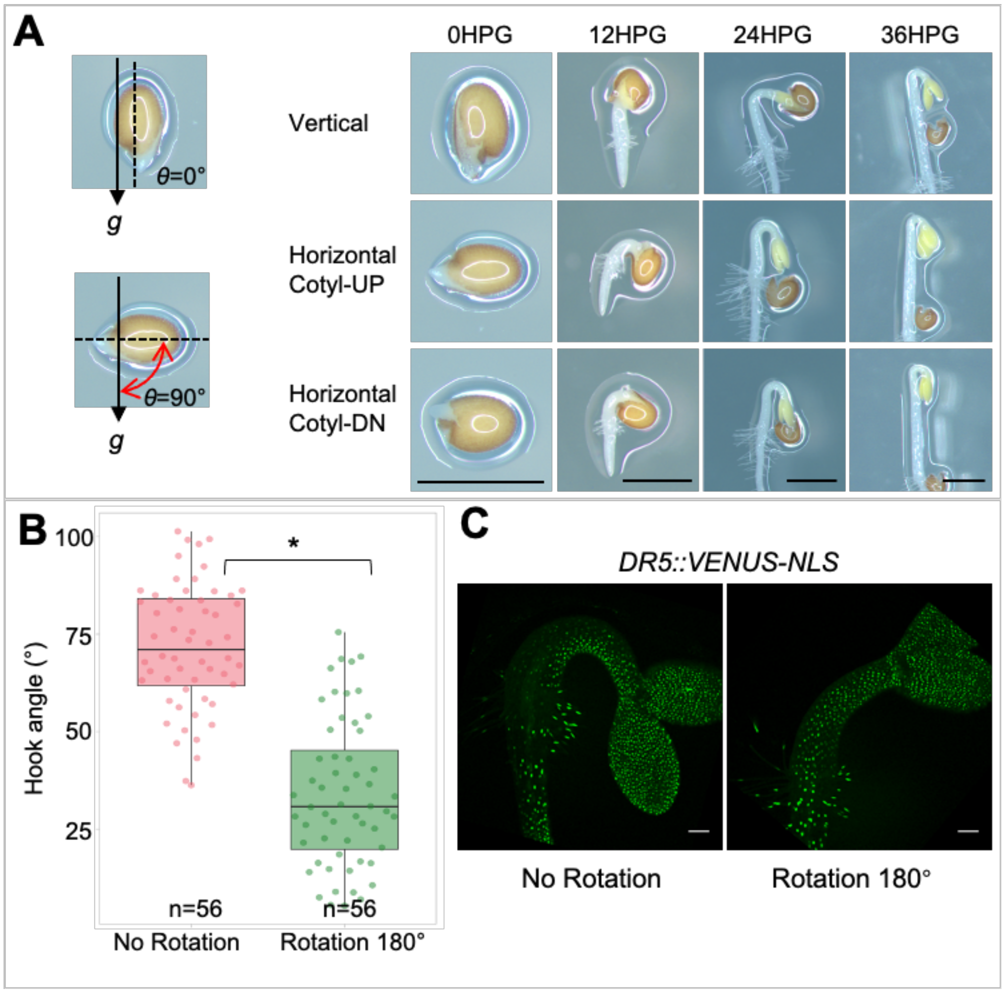


**Fig. S4. Gravity is critical for the formation of apical hook.**

**(A)** Schematic representation of the inclination angle $\theta$ relative to the gravity vector. Images of the apical hook development in differentially oriented seedlings. Scale bars=1mm. **(B)** Quantification of angles of apical hook in the periodically rotated seedlings. **(C)** *DR5::VENUS-NLS* expression in the apical hook in the periodically rotated seedlings. Scale bars=100 µm. For **(B** and **C)**, germinated seeds were initially placed at a horizontal orientation at 0HPG, the seedlings were then rotated by 0 or 180 degrees once per hour. Images were acquired and angles of the apical hook were measured at 12HPG, n=56 hooks from three replicates. * indicates $P$ value < 0.01.

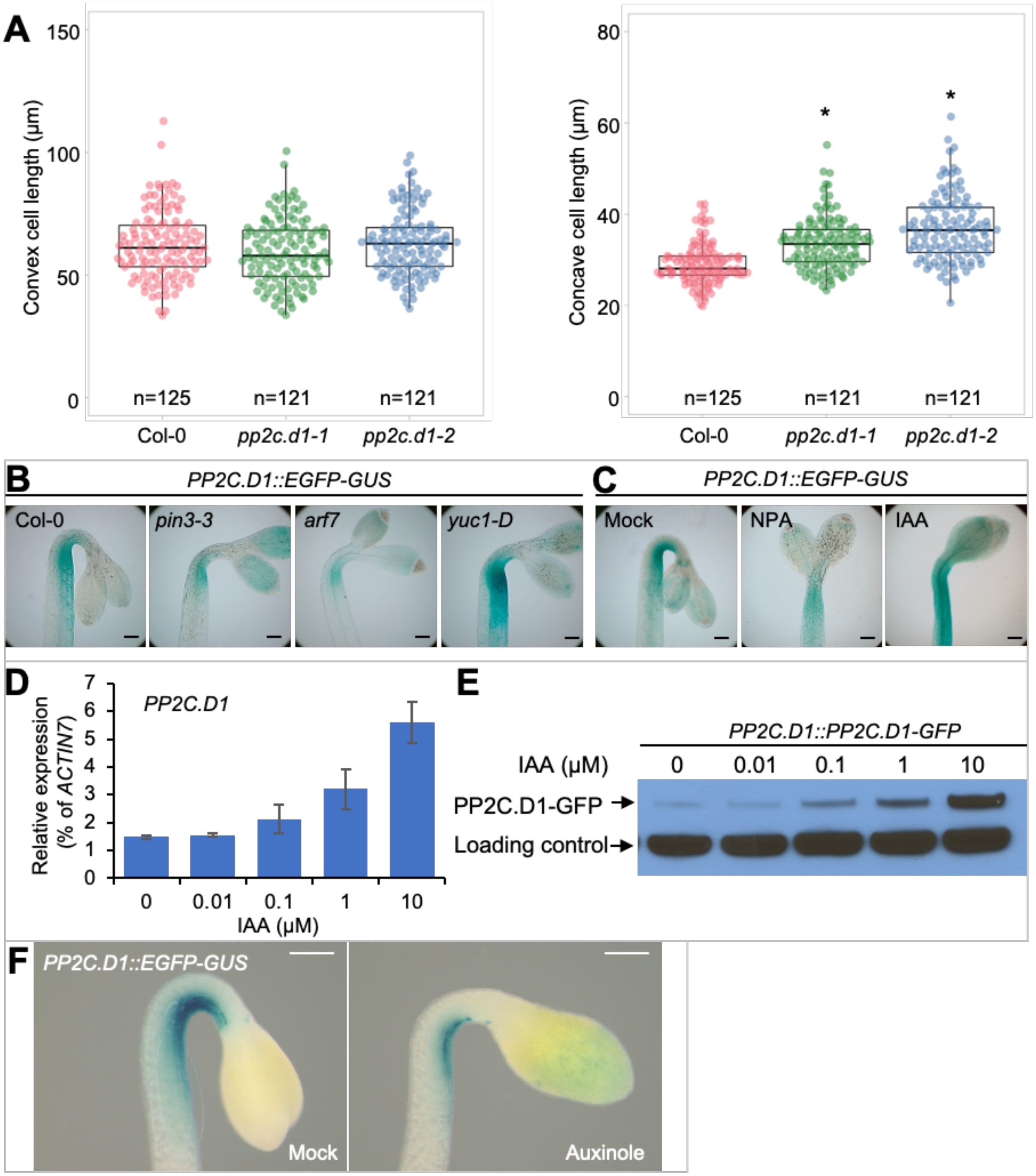


**Fig. S5. Auxin-induced *PP2C.D1* inhibits cell elongation at the concave side of the apical hook.**

**(A)** Quantification of epidermal cell length at the concave and convex sides of the apical hook in Col-0, *pp2c.d1-1*, and *pp2c.d1-2*. * indicates *P* value < 0.001. **(B)** *PP2C.D1::EGFP-GUS* expression in Col-0, *pin3-3*, *arf7* and *yuc1-D* at 36 HPG. Scale bars=100 μm. **(C)** *PP2C.D1::EGFP-GUS* expression when grown on media containing 1 μM NPA or 10 μM IAA at 36 HPG. Scale bars=100 μm. **(D)** RT-qPCR analysis showing that auxin induces *PP2C.D1* expression in a dose-dependent manner. *PP2C.D1* transcript levels were normalized against *ACTIN7* expression. Data are means ± s.e.m. from three replicates. **(E)** Western blot showing that auxin induces *PP2C.D1::PP2C.D1-GFP* expression in a dose-dependent manner. A non-specific band is shown as a loading control. **(F)** *PP2C.D1::EGFP-GUS* expression after auxinole treatment. *PP2C.D1::EGFP-GUS* seedlings grown on media supplemented with DMSO (mock) or 10 μM auxinole prior to GUS staining at 36HPG. Scale bars=200 μm.

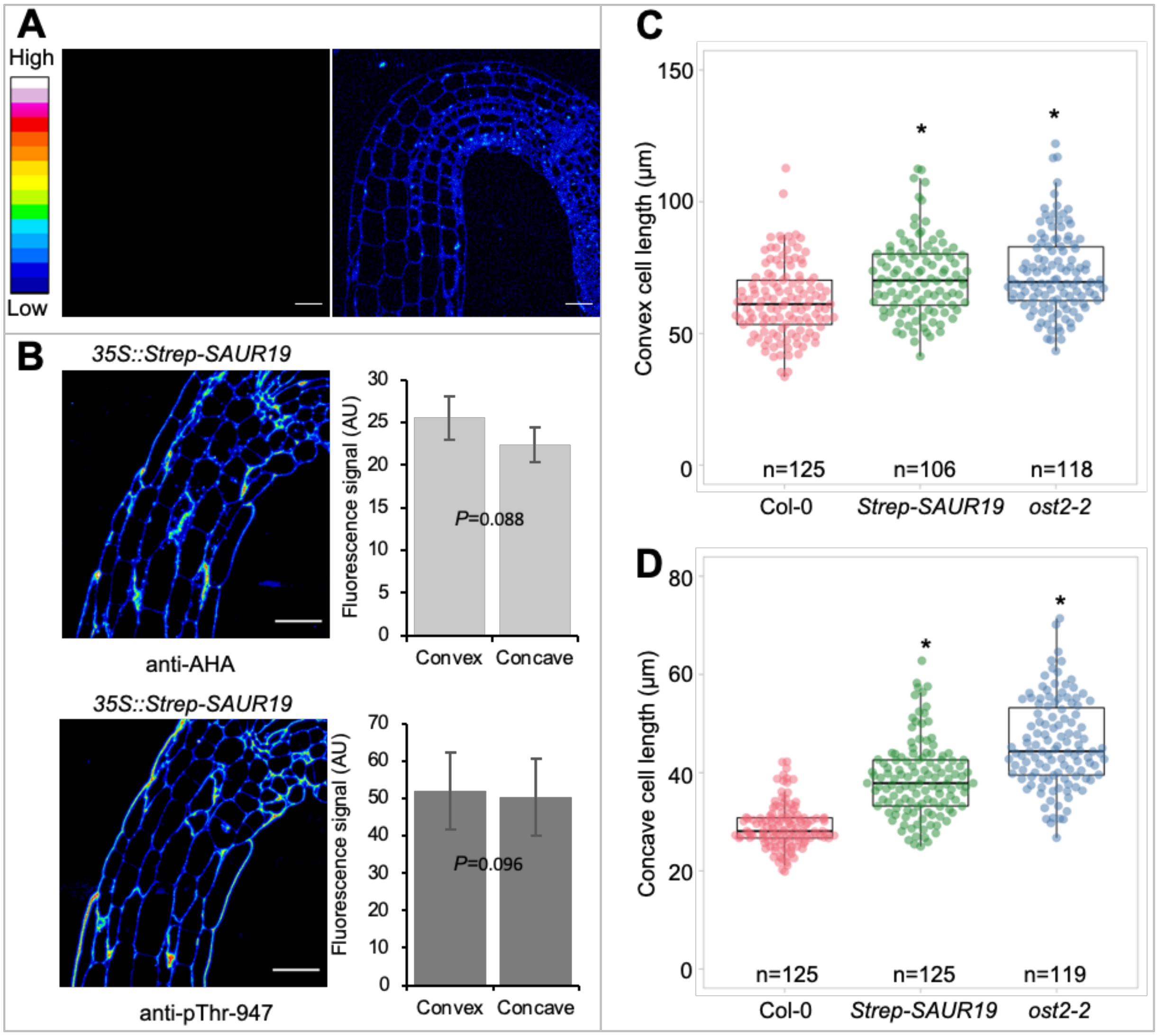


**Fig. S6. Asymmetric acid growth is critical for apical hook development. (A)** Negative control micrographs for immunolabelling of a wild type hypocotyl during the hook maintenance stage: PBS with 2% BSA was used instead of the primary antibody and all the other steps were kept the same (left). The right image is the same image as the left with the gain extremely increased to show the tissue outline. Scale bars=50 µm. **(B)** Immunolabeling and signal quantification of PM $H^+$-ATPase and $Thr^{947}$-phosphorylated PM $H^+$-ATPase during apical hook development in the *35S::Strep-SAUR19* seedlings. Scale bars=50 µm. Details of the signal quantification can be found in the methods section. **(C** and **D)** Quantification of epidermal cell lengths on the convex and concave sides of the apical hook in Col-0, *35S::Strep-SAUR19* and *ost2-2*. * indicates $P$ value < 0.001.

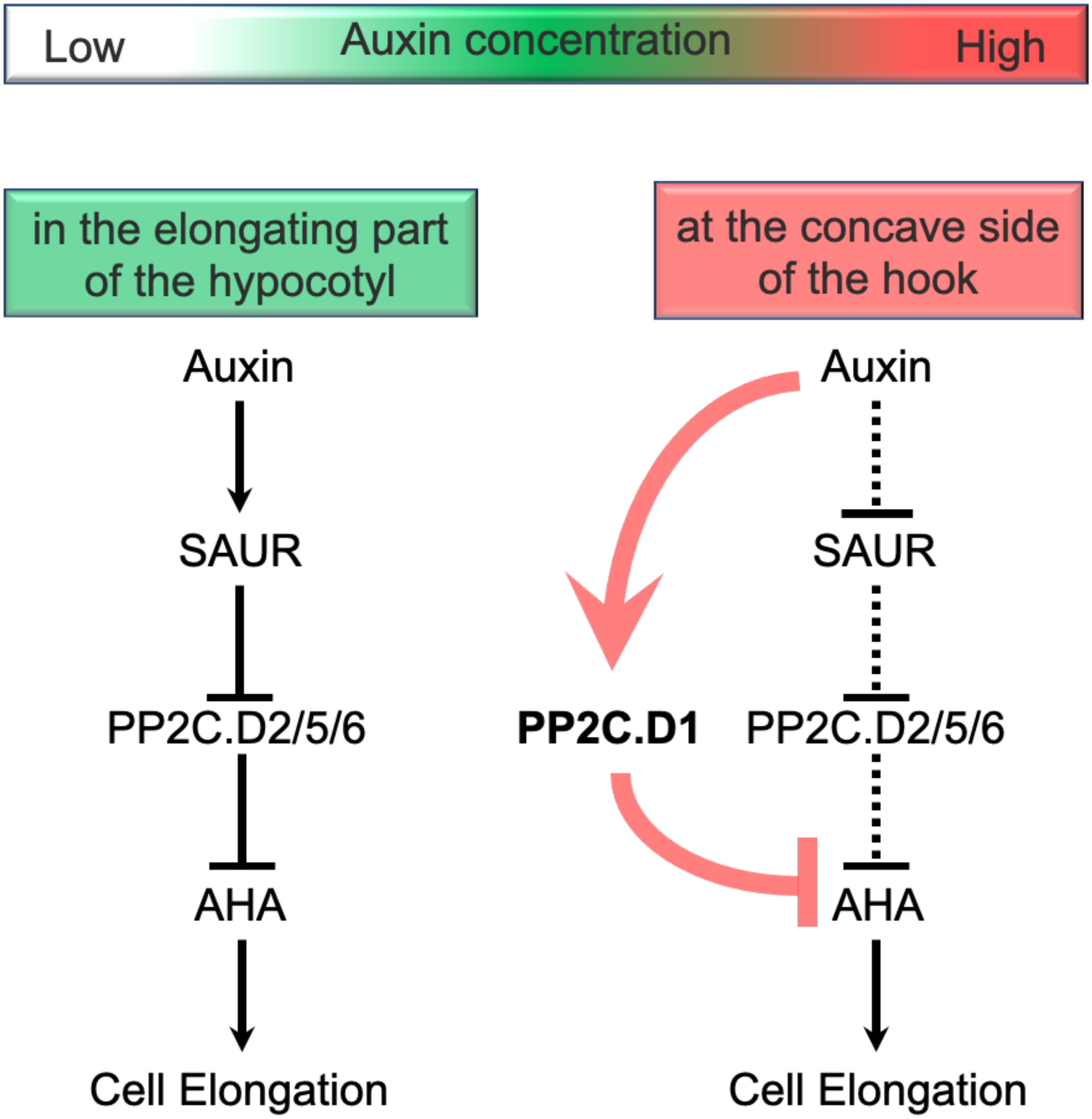


**Fig. S7. Proposed model for auxin-mediated regulation of cell elongation during etiolated seedling development.** In the elongating part of the hypocotyl, auxin promotes cell elongation through an acid-growth mechanism, in which auxin-induced SAURs inhibit PP2C.D2/5/6 phosphatases to activate the PM $H^+$-ATPase. When auxin concentrations in the cells are high (e.g. in epidermal cells of the concave side of the hook), *PP2C.D1* expression is induced. This enables auxin to bypass SAUR-regulation to directly activate *PP2C.D1* for inhibiting PM $H^+$-ATPase activity and cell elongation. In addition, high auxin levels may repress expression of some *SAUR* genes, providing another way for auxin to inhibit PM $H^+$-ATPase activity and cell elongation. The inhibition bars between auxin-SAUR-PP2C.D2/5/6-AHA are dashed, as how high auxin levels on the concave side of the hook affect SAUR-PP2C.D2/5/6 modules is uncertain.

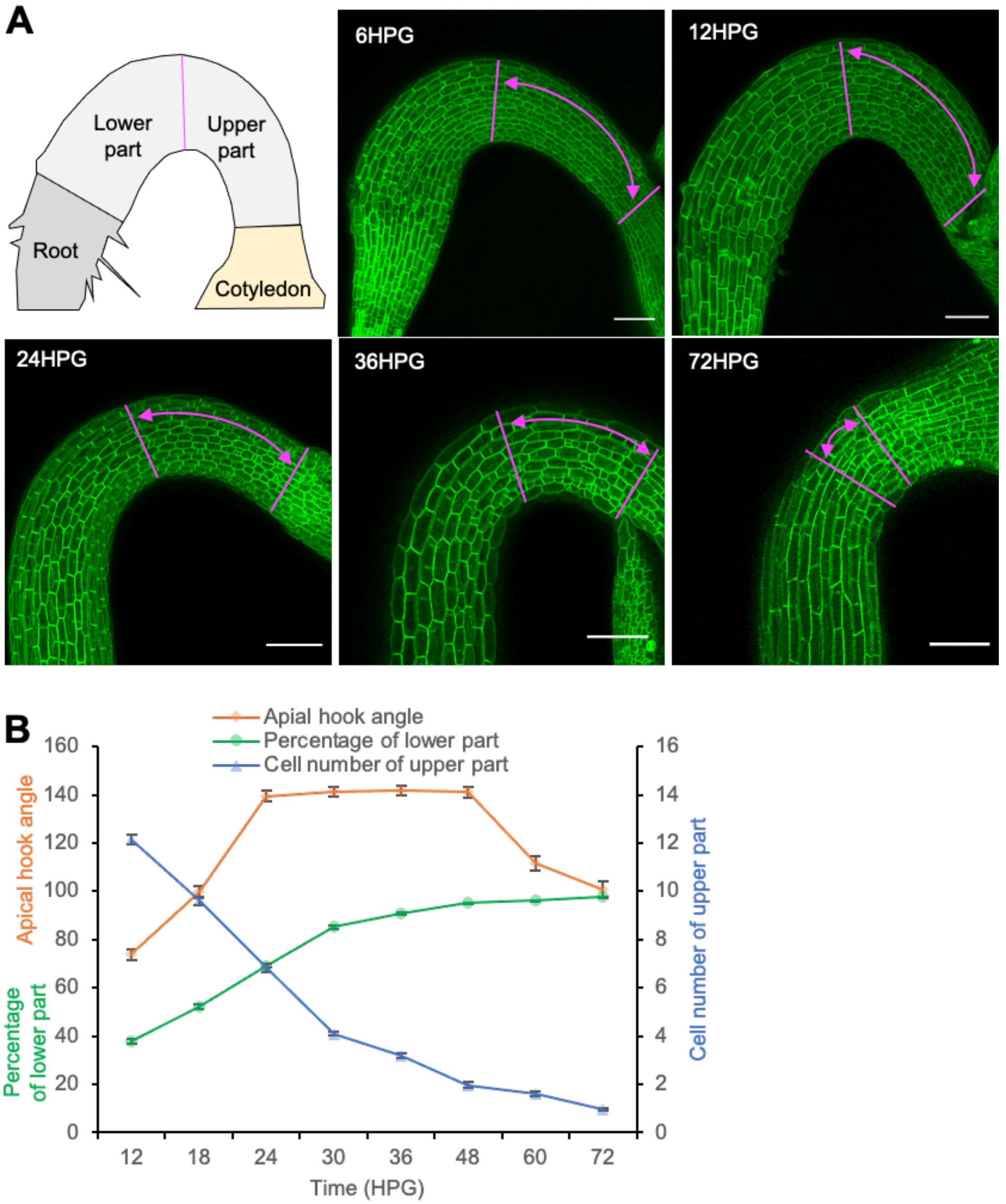


**Fig. S8. The apical hook moves upwards during hypocotyl development. (A)** Apical hook micrographs for the indicated times. Etiolated MYR-YFP seedlings were used for confocal imaging. Magenta lines were used to mark the tip of the hook for separating the lower and the upper parts of the hypocotyls. The tip of the hook was determined using the kappa plugin in FIJI. Briefly, the points with highest curvature on the concave and convex sides were determined and a line joining them indicates the middle of the hook. Schematic representation of etiolated Arabidopsis seedling was shown to define the lower and upper parts of the hypocotyl. Scale bars=100 µm. **(B)** Quantification of the kinetics of percentages of the lower part of the hypocotyl (n=32) and cell numbers of the upper part (n=30) during apical hook development. Values represent sample means ± s.e.m. from three replicates. Germinated seeds were initially placed at a horizontal orientation at 0HPG.

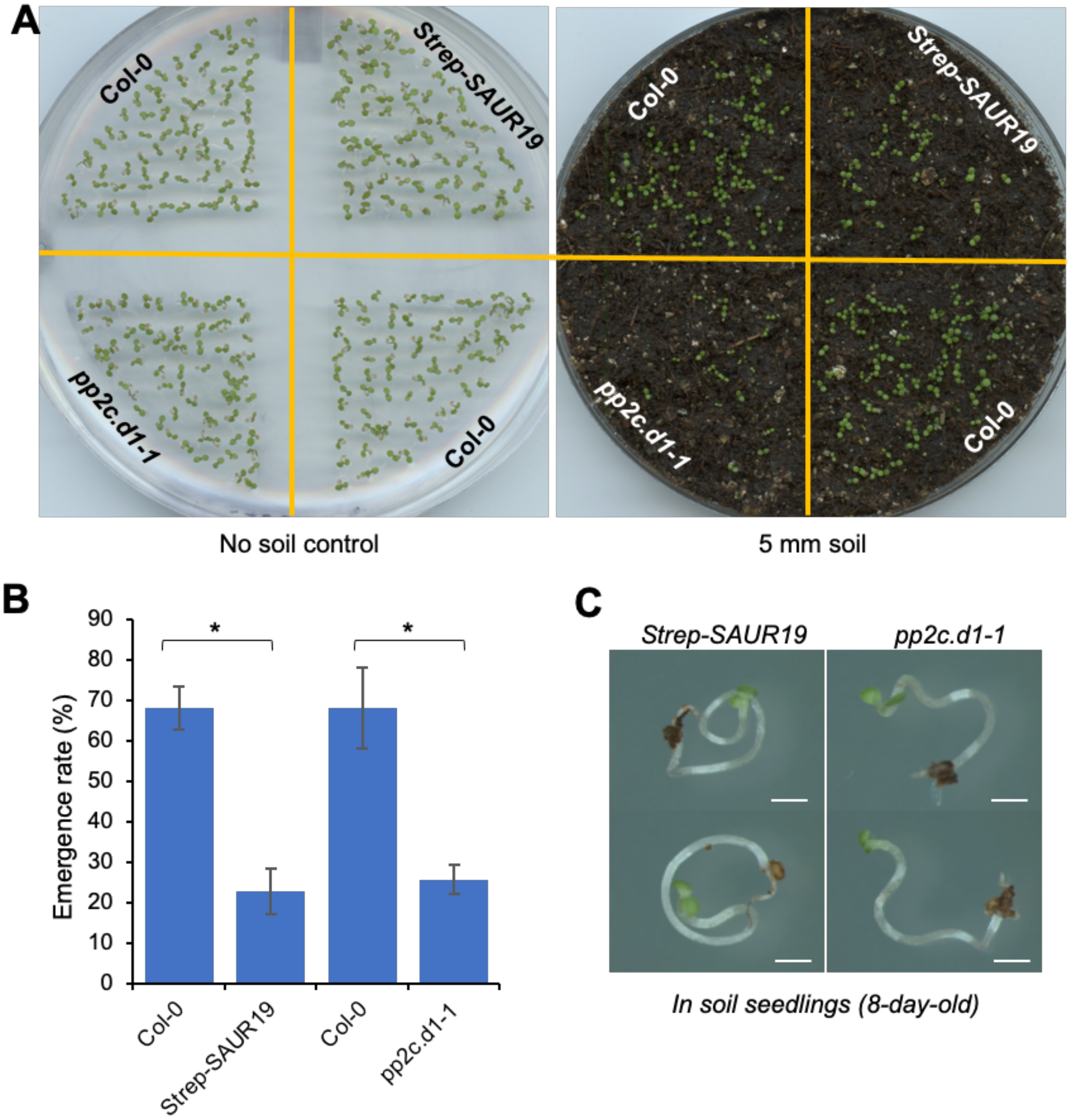


**Fig. S9. Proper etiolated development is critical for seedling emergence.** Seedling emergence phenotypes (**A**) and quantitative analysis (**B**) of Col-0, *35S::Strep-SAUR19* and *pp2c.d1-1* mutant. Germinated seeds were covered with a 5-mm layer of soil, and then grown under continuous white light. Pictures were acquired and the number of emerged seedlings were counted at 60HPG. * indicates *P* value < 0.05. **c**, Phenotypes of un-emerged *35S::Strep-SAUR19* and *pp2c.d1-1* seedlings after extricating them from the soil. Scale bars=1 mm.

**Table S1 | Primers used in this study.**

| Purpose | Name | Sequence (5'-3') | Note |
|---|---|---|---|
| ChIP-qPCR | PP2C.D1-P1-F | ggaagttgtgtaacccacttgttg | PP2C.D1 promoter regions |
| ChIP-qPCR | PP2C.D1-P1-R | gctactgtagagagaaccactgtcg | PP2C.D1 promoter regions |
| ChIP-qPCR | PP2C.D1-P2-F | ccgtttattatatttgtggagcct | PP2C.D1 promoter regions |
| ChIP-qPCR | PP2C.D1-P2-R | tcttctagctcacgagtcac | PP2C.D1 promoter regions |
| ChIP-qPCR | PP2C.D1-P3-F | cgacagtggttctctctacagtagc | PP2C.D1 promoter regions |
| ChIP-qPCR | PP2C.D1-P3-R | cagatcgaaattcatgtcatgg | PP2C.D1 promoter regions |
| ChIP-qPCR | PP2C.D1-P4-F | ctcacaacaagtaagagatcagc | PP2C.D1 promoter regions |
| ChIP-qPCR | PP2C.D1-P4-R | gatttgagagctgagaaagactg | PP2C.D1 promoter regions |
| ChIP-qPCR | PP2C.D1-P5-F | tggttccaagttcaatgctccc | PP2C.D1 promoter regions |
| ChIP-qPCR | PP2C.D1-P5-R | ggtaacaatatctcatgtaaacgtctg | PP2C.D1 promoter regions |
| ChIP-qPCR | PP2C.D1-P6-F | cagacgtttacatgagatattgttacc | PP2C.D1 promoter regions |
| ChIP-qPCR | PP2C.D1-P6-R | ctcctctttactgttcgatgatg | PP2C.D1 promoter regions |
| ChIP-qPCR | PP2C.D1-P7-F | catcatcgaacagtaaagaggag | PP2C.D1 promoter regions |
| ChIP-qPCR | PP2C.D1-P7-R | ggtgttggaaagcagggaaagaggcc | PP2C.D1 promoter regions |
| ChIP-qPCR | PP2C.D1-P8-F | cacacttactctagggcttccac | PP2C.D1 promoter regions |
| ChIP-qPCR | PP2C.D1-P8-R | ctttctcaggtatgtaatcttcgagc | PP2C.D1 promoter regions |
| RT-qPCR | PP2C.D1-F | AATGGCCTACGAACCCACAG | |
| RT-qPCR | PP2C.D1-R | AGCGGTGCTTCATCACAAGA | |
| RT-qPCR | ACTIN7-F | CCATTCAGGCCGTTCTTTC | |
| RT-qPCR | ACTIN7-R | CGTTCTGCGGTAGTGGTGA | |
| RT-qPCR | SAUR22-F | GACAAATAGAGAATTATAAATGGCTCTG | |
| RT-qPCR | SAUR22-R | ATGAATTAAGTCTATATCTAACTCGGAAA | |
| RT-qPCR | SAUR19-F | GATTCTAAGCCGCTCCAC | |
| RT-qPCR | SAUR19-R | CCGAGAAGTCACATTGATG | |
| RT-qPCR | SAUR9-F | TCAACACCGAAGTCGCTATG | |
| RT-qPCR | SAUR9-R | TCGTGCTCGAAACCAAACTC | |
| RT-qPCR | IAA2-F | TTACGGGAAGATCTCACTGG | |
| RT-qPCR | IAA2-R | ATCCAAAAGCAATGGCGTAC | |
| RT-qPCR | IAA5-F | CGGCGAAAAAGAGTCAAGTTGTGGGT | |
| RT-qPCR | IAA5-R | CATTCACTTTCCTTCAACGTATCATCA | |
| RT-qPCR | GH3.3-F | ATCACAGAGTTCCTCACAAGC | |
| RT-qPCR | GH3.3-R | TTGCCTTTGTCTAATCCGGG | |

**Movie S1. Apical hooks reorient to the gravity direction.** Apical hook development was recorded with an infrared light source by a spectrum-enhanced camera. Upon turning seedlings to a horizontal plane at 36HPG, apical hooks tend to actively reorient to the gravity vector. The magenta and white arrowheads indicate hooks, for which cotyledons were either above or below the hypocotyl just after the turning at 36HPG. Scale bar = 5 mm.

**Movie S2. Apical hook development in differentially orientated seedlings.** Apical hook development of differentially orientated seedlings was recorded with an infrared light source by a spectrum-enhanced camera. Germinated Col-0 seeds were initially placed at vertical or horizontal orientations at 0HPG.

**Movie S3. Apical hook development in WT and *shr-2* seedlings.** Apical hook development of WT and *shr-2* seedlings was recorded with an infrared light source by a spectrum-enhanced camera. Germinated seeds were initially placed at a horizontal orientation at 0HPG.

**Movie S4. Apical hook development in WT and *atlazy1,2,3,4* seedlings.** Apical hook development of WT and *atlazy1,2,3,4* seedlings was recorded with an infrared light source by a spectrum-enhanced camera. Germinated seeds were initially placed at a horizontal orientation at 0HPG.
